# Dynamical nonequilibrium molecular dynamics simulations identify allosteric sites and positions associated with drug resistance in the SARS-CoV-2 main protease

**DOI:** 10.1101/2022.12.10.519730

**Authors:** H. T. Henry Chan, A. Sofia F. Oliveira, Christopher J. Schofield, Adrian J. Mulholland, Fernanda Duarte

## Abstract

The SARS-CoV-2 main protease (M^pro^) plays an essential role in the coronavirus lifecycle by catalysing hydrolysis of the viral polyproteins at specific sites. M^pro^ is the target of drugs, such as nirmatrelvir, though resistant mutants have emerged that threaten drug efficacy. Despite its importance, questions remain on the mechanism of how M^pro^ binds its substrates. Here, we apply dynamical nonequilibrium molecular dynamics (D-NEMD) simulations to evaluate structural and dynamical responses of M^pro^ to the presence and absence of a substrate. The results highlight communication between the M^pro^ dimer subunits and identify networks, including some far from the active site, that link the active site with a known allosteric inhibition site, or which are associated with nirmatrelvir resistance. They imply that some mutations enable resistance by altering the allosteric behaviour of M^pro^. More generally, the results show the utility of the D-NEMD technique for identifying functionally relevant allosteric sites and networks including those relevant to resistance.

## Introduction

The ongoing COVID-19 pandemic has led to worldwide efforts to develop antiviral drugs effective against the SARS-CoV-2 coronavirus. The SARS-CoV-2 main protease (M^pro^) cleaves viral polyproteins into functional units essential for viral replication and pathogenesis and is a validated drug target.^1^ M^pro^ hydrolyses the amide bond between P1 and P1′ residues in the conserved sequence: [P4=Ala/Val/Pro/Thr]-[P3=X]-[P2=Leu/Phe/Val]-[P1=Gln]-[P1′=Ser/Ala/Asn], where X is any residue.^2-4^ Due to the essential role of M^pro^, the apparent absence of human proteases with a similar substrate selectivity, and the high sequence identity (96%) between SARS-CoV and SARS-CoV-2 M^pro^s, inhibitors targeting M^pro^ enzymes are likely useful for treatment of many coronavirus diseases.^5-8^ The covalently acting M^pro^ inhibitor PF-07321332 (nirmatrelvir), the active ingredient in Paxlovid, is the first oral COVID-19 drug that has been authorised for emergency use by the FDA.^9, 10^ Another M^pro^ inhibitor in clinical-stage development is S-217622 (ensitrelvir), a noncovalently binding, nonpeptidic inhibitor that was discovered following virtual screening and medicinal chemistry optimisation.^11^

Crystal structures^4, 12-14^ and molecular simulations^15-20^ have provided insights into M^pro^ structure, dynamics and interactions with its substrates and other ligands. Molecular simulations of M^pro^ have identified cryptic pockets,^21^ informed on mechanisms of inhibition,^22-24^ and helped to suggest potential lead compounds for M^pro^ inhibitor design.^25, 26^ We have identified conserved interactions between M^pro^ and its 11 native substrates, each modelled as an 11-mer P6-P5′ peptide.^20^ However, how the binding of substrates or inhibitors induces changes in the M^pro^ structure and its dynamics has been unclear. Changes in dynamics and conformational behaviour have been shown to be the cause of mutation-induced drug resistance for some targets.^27-30^ For instance, mutations distal to the active site of the HIV-1 protease have been shown to confer resistance to darunavir through alterations in conformational dynamics.^31^ Given the relatively rapid rate at which SARS-CoV-2 mutates, M^pro^ targeting drug resistance is a major concern.^8, 32, 33^ A detailed understanding of how M^pro^ recognises its natural substrates thus may aid the development of next-generation inhibitors.^12, 34^

Here, we describe the application of dynamical-nonequilibrium molecular dynamics (D-NEMD)^35-38^ simulations to investigate the response of M^pro^ to the instantaneous removal of a substrate in one of the active sites in the dimer. The D-NEMD approach aims to identify in the protein any structural changes, which refer to conformational rearrangements, as well as dynamic changes, which relate to changes in fluctuations, following a perturbation. D-NEMD simulations are emerging as a useful tool to probe signal propagation and allosteric effects,^38^ *e*.*g*., they have identified a general mechanism of signal propagation in nicotinic acetylcholine receptors;^39, 40^ communication pathways between allosteric and active sites in clinically relevant β-lactamases;^30^ and modulation of SARS-CoV-2 spike protein behaviour by pH^41^ and ligand occupancy in the fatty acid binding site.^42-44^ Notably in this context, the pathways identified in the β-lactamases contain positions that differ between enzymes with different spectrums of antibiotic breakdown activity.^30^

In the D-NEMD approach, the system is sampled first by equilibrium molecular dynamics (MD) simulations. Configurations from these trajectories provide starting points for multiple short nonequilibrium simulations, through which the effect of a perturbation can be determined using the Kubo-Onsager relation.^37, 38^ Running a large number (usually tens to hundreds) of nonequilibrium simulations allows the statistical significance of the response of the protein to the perturbation to be tested.^37^

The results here identify structural pathways involved in M^pro^ communication, both within and between subunits of the dimer. We use a substrate sequence representing the nsp8/9 P6-P5′ substrate (Ser-Ala-Val-Lys-Leu-Gln|Asn-Asn-Glu-Leu-Ser, where “|” denotes the cleavage site), which we have previously characterised by experiment and simulation, and refer to as s05 according to previous nomenclature.^20^ s05 was selected because it is a conserved substrate in coronaviruses,^4^ and its P4-P1 sequence is reported to be optimal for M^pro^ activity amongst natural amino acid sequences.^1^

## Results and Discussion

### 1. Equilibrium simulations of the M^pro^-substrate complex

Initially, five independent equilibrium MD simulations of 200 ns (an aggregate of 1 μs) were carried out for homodimeric M^pro^ noncovalently complexed with a single 11-mer peptide substrate s05 (**Figure 1a, b**), using established protocols.^20^ s05 occupies the active site formed predominantly by residues of the M^pro^ chain which is denoted as chain A (ChA), while the other (chain B, ChB) active site was simulated without a substrate. In subsequent descriptions, all residues relate to M^pro^ chain A (Ser1-Gln306), except chain B residues which are indicated by prime (Ser1′-Gln306′).

**Figure 1:**
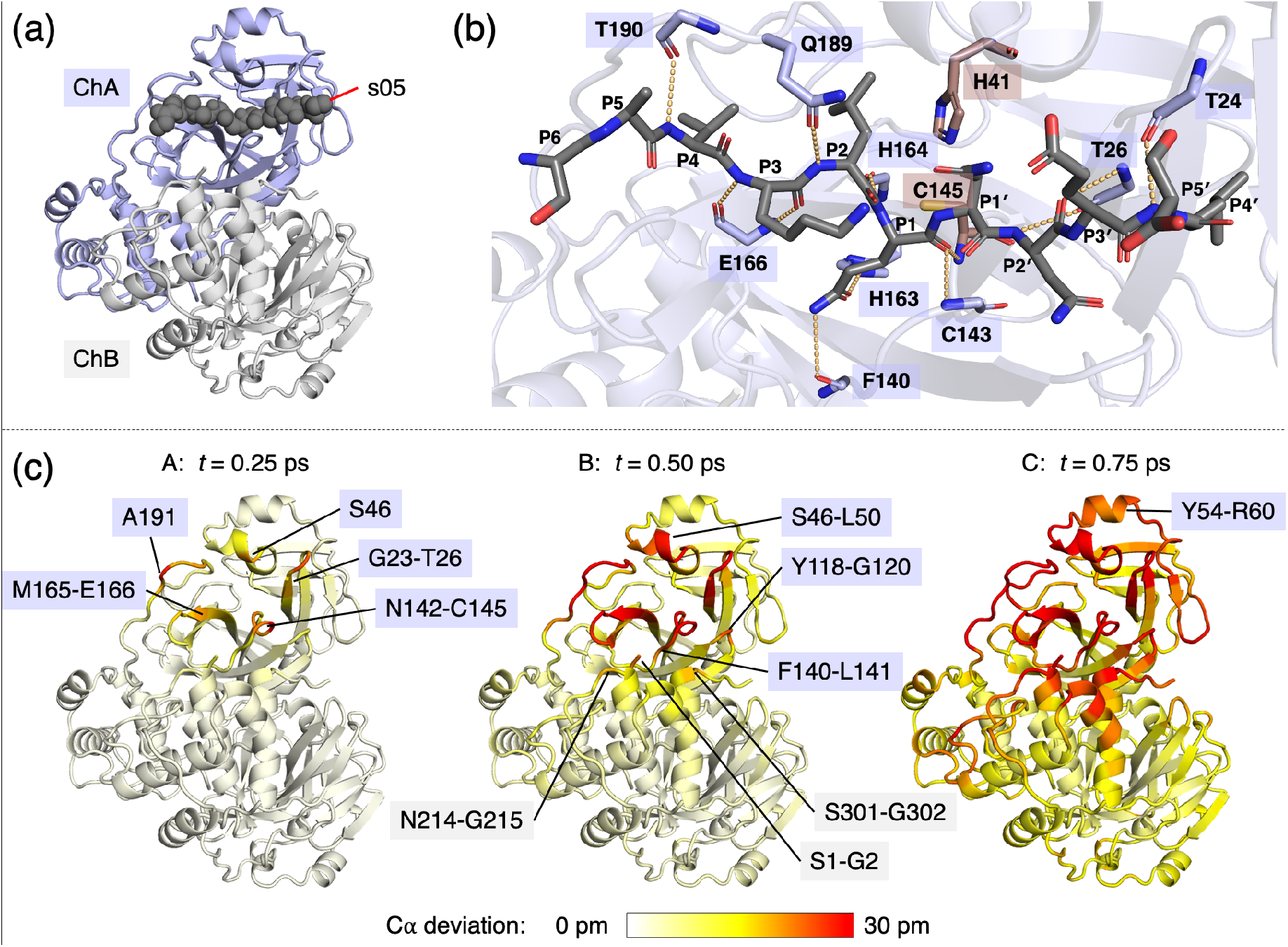
The immediate response of M^pro^ to substrate removal from the equilibrated M^pro^-substrate complex. (a) View of the M^pro^ dimer (chains A and B shown as blue and grey cartoons, respectively) complexed with the s05 peptide substrate (backbone shown as black spheres) prior to MD simulations. (b) View of a MD-derived snapshot, with the 12 conserved HBs (**Figure S1.9**) shown as orange dotted lines. s05, the M^pro^ catalytic dyad, and other HB-forming M^pro^ residues are shown as black, brown, and blue sticks respectively (hydrogens omitted). (c) The initial sub-ps response of M^pro^ to s05 removal in terms of Cα deviation represented on a white-yellow-red scale. All figures were generated using PyMOL.^45^

Analysis of RMSD and RMSF values, protein secondary structure content, and M^pro^-substrate interactions confirm the stability of the M^pro^ dimer-s05 complex, with the system considered to be well equilibrated from *t* = 50 ns onwards in all simulations (**SI Section S1**). The 12 M^pro^-substrate hydrogen bonds (HBs) previously found to be conserved across 11 native substrates are also observed here (**Figures 1b, S1.9-11**).^20^ These equilibrium trajectories were then used as the basis for D-NEMD simulations.

### 2. Immediate effect of substrate removal

To analyse the response of the M^pro^ dimer to substrate removal, snapshots were extracted every 5 ns from *t* = 50 to 195 ns from all five equilibrium simulations (a total of 150 conformations), and the s05 peptide instantaneously annihilated (see **Methods**). The time-dependent response of the protein to the annihilation of s05 was monitored using the Kubo-Onsager relation,^37, 38^ by subtracting the position of each Cα atom between the nonequilibrium and equilibrium trajectories at equivalent time points (**Figure S2.1)**. Cα movements were used, employing the same approach as in previous investigations of other systems, as they are less susceptible to thermal fluctuations than those of sidechain atoms, and they normally reflect the most substantial responses in the protein.^38^ The subtraction approach enables a clear identification of the response via cancellation of most random fluctuations between the equilibrium and nonequilibrium trajectories.^30, 39^ Note that the perturbation (in this case, the deletion of the s05 peptide) is not intended to simulate the physical process of substrate binding or dissociation, but rather to trigger a fast response from M^pro^ as it adapts to s05 removal. Such a perturbation forces the system out of equilibrium, thus creating the driving force for conformational changes to occur. Similarly, the simulation timescale does not reflect the real timescale of conformational changes.^39^

Soon after the perturbation, a clear response from M^pro^ is observed, notably in residues directly involved in substrate binding (**Figures 1cA, S2.2; Table S2.1**). The statistical significance of the response is confirmed by determination of standard errors of the mean (SEM) (**Figure S2.3**), as performed in previous investigations of other systems.^30, 42^ At *t* = 0.25 ps, the most perturbed regions include: Gly23-Thr26, which forms HBs with the substrate P2′ and P4′ residues (**Figure 1b**); Asn142-Cys145, which constitutes the oxyanion hole loop that stabilises the scissile amide carbonyl oxygen; Met165, which forms the S2 hydrophobic pocket; Glu166, whose backbone forms two strong HBs with the substrate P3 backbone; and Ala191, which is adjacent to Thr190 that binds the substrate P4 residue via a HB. Hence, as expected, the residues with the largest initial deviations are those originally in direct contact with, or very close to, the substrate, and these residues are likely to rearrange to accommodate the substrate.

Notable initial responses are also observed in residues belonging to chain B (**Figures 1cB, S2.2-3**). Chain B residues which show a deviation >15 pm at *t* = 0.50 ps include: Ser1′, Gly2′, Asn214′, Gly215′, Ser301′ and Gly302′. These residues are in the vicinity of chain A regions that also show significant responses, such as Phe140-Leu141 and Tyr118-Gly120 (**Figure S2.4**). These responses highlight the importance of inter-subunit dynamics, where structural and dynamic changes occurring in one are swiftly propagated to the other. For example, the structural importance of the N-terminus (Ser1′-Gly2′) of chain B is consistent with its role of forming part of the S1 pocket that accommodates the conserved P1 Gln substrate residue.^5, 6^ Asn214′ and Gly215′ are further away from the substrate binding site; notably, the N214A variant of SARS-CoV M^pro^ is much less active than the wild type enzyme against a fluorogenic substrate peptide, despite being similar in dimerization affinity and structure, with the inactivation proposed to be driven by changes in protein dynamics.^15, 46^ Finally, the observed perturbations in Ser301′ and Gly302′ of the C-terminal tail of chain B are consistent with crystallographic observations that this region adopts distinct conformations upon the binding of substrate or the peptidomimetic inhibitor N3.^13, 47^ Overall, the D-NEMD analysis highlights the communication between chains A and B and, in agreement with reported experimental and crystallographic data, identifies three structurally and dynamically important regions of chain B.

At 0.75 ps, the structural responses intensify and propagate further from the substrate binding site into both chains A and B (**Figures 1cC, S2.3**). These responses include signal transmission in chain A from Ser46 to the whole Ser46-Leu50 helix, and then to the Tyr54-Arg60 helix, above the active site. There is also transmission through the helical regions in chain B away from the dimer interface. These results show communication between the subunits of the dimer.

### 3. Propagation of signal to allosteric sites

The conformational changes at the dimer interface are of particular interest. While the initial response (**Figure 1c**) in chain B is concentrated at dimer interface regions with little secondary structure, such as Asn214′-Gly215′ and Ser301′-Gly302′, notable response in the Gln244′-Thr257′ kinked helix is observed across the ps timescale (**Figure 2**). The helix residues Pro252′, Leu253′, and Gln256′; the C-terminal tail residues Cys300′, Ser301′, and Gly302′; Ile213′, which is adjacent to the perturbed Asn214′-Gly215′ region; and the earlier highlighted chain A residues Tyr118, Leu141, and Asn142 all form an allosteric site (coloured orange in **Figure 2a**), which binds small molecules as demonstrated by crystallography.^7, 48, 49^ One such compound is pelitinib, which has high antiviral activity and which does not bind at the active site.^48^ It is striking that the D-NEMD simulations, which were initiated by substrate removal, identify this allosteric site. Our results show a direct connection between the allosteric site and the active site via conformational motions. Based on these D-NEMD results, it may be proposed that the presence of a ligand at this allosteric site will modulate enzyme dynamics, including those affecting the rate and/or affinity of binding of substrates. Such a ligand may also alter how dynamic and structural changes are communicated within the protein, a likely factor in allosteric inhibition.

**Figure 2:**
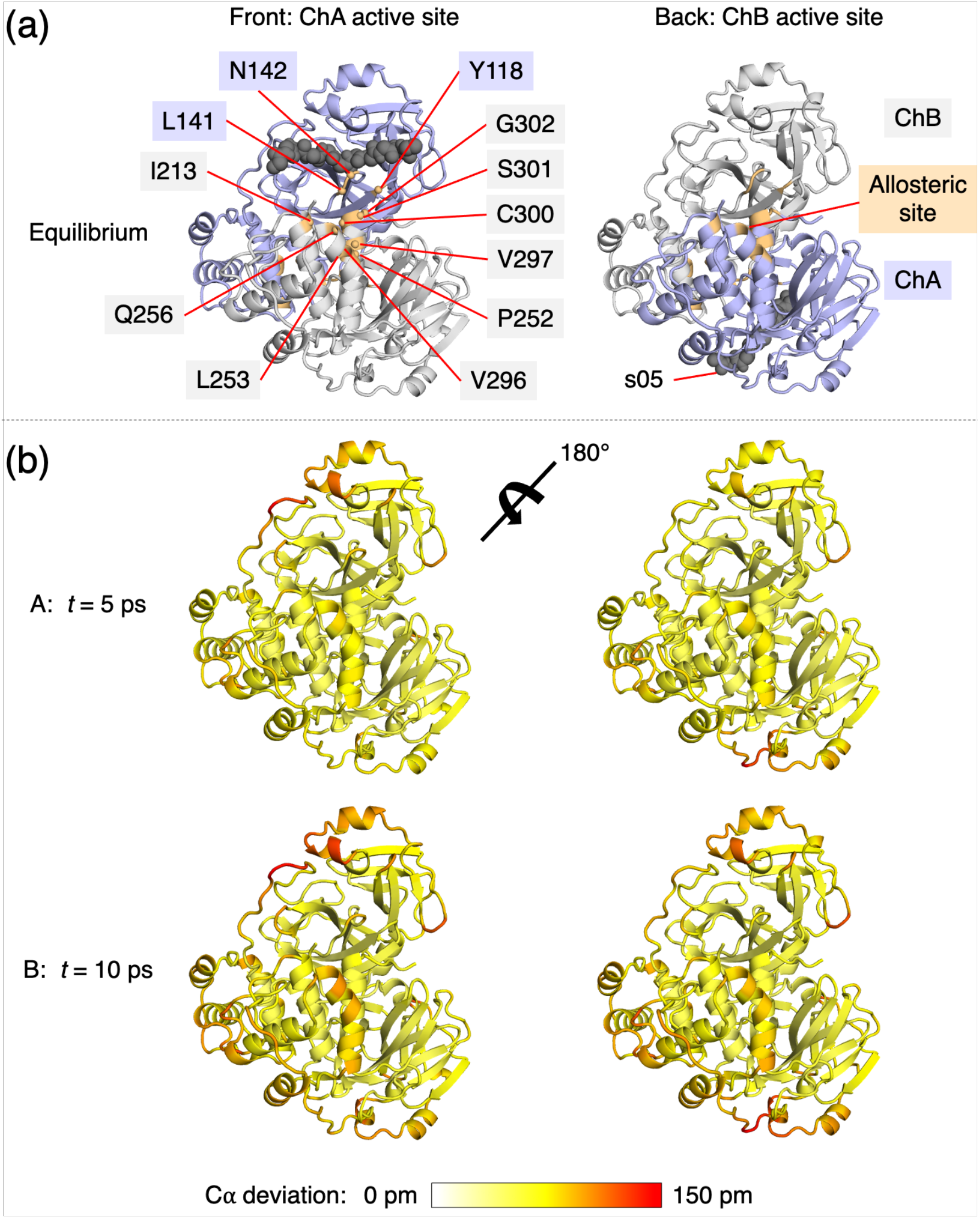
D-NEMD identifies connection between the active site and a known allosteric inhibition site. (a) Alternative (180° rotated) views of the M^pro^-s05 complex prior to MD simulations. Residues constituting the pelitinib binding allosteric site (within 4 Å of pelitinib according to PDB 7AXM: Tyr118, Leu141, Asn142, Ile213′, Pro252′, Leu253′, Gln256′, Val296′, Val297′, Cys300′, Ser301′, Gly302′; see **Figure S3.6**)^48^ are labelled and shown in orange. (b) D-NEMD Cα responses to s05 peptide removal shown on a white-yellow-red scale, with red indicating large deviation. The figures were generated using PyMOL.^45^

As the time following substrate removal increases across the ps and ns timescales, the structural response in residues in terms of Cα deviations increases and spreads across the M^pro^ dimer (**Section S3**). Perturbed allosteric regions are found in both M^pro^ chains, and except for the region around Gln189-Ala191, which has higher deviation in chain A than B, the deviations in the two chains are similar (**Figures S3.2-4**). Some of the most perturbed residues (*e*.*g*., Asn277 and Gly278 in both chains at *t* = 1 ns) also display high mobility in the equilibrium simulations (**Figure S1.3**). This is unsurprising, because more flexible regions require less energy to change their positions.^36, 50^

### 4. Coherent displacements in residues upon substrate removal

Complementarily to the above approach, which averages scalar deviations across replicas, the displacement vector of Cα atoms between the equilibrium and nonequilibrium trajectories at equivalent time points can be averaged. The resulting signals indicate the average direction of the response of the residues upon the perturbation; however, it is important to note that the responses of residues that are dynamically affected in multiple directions or in a disordered manner are not apparent in this analysis.

The response of M^pro^ to substrate removal calculated from Cα displacement vectors is shown in **Figure S4.1**, with the statistical significance of the responses assessed by SEM (**Figures S4.2-5**). Due to the cancellation of random movements in mobile atoms, the observed responses are more localised than those calculated by averaging scalar deviations (**Figures S3.1, S4.1**). Upon removal of the substrate peptide, M^pro^ regions that form the substrate binding site show responses with well-defined directions (**Figure 3aA; Table 1**). These regions include: Thr26-Leu27 in the S1′-S2′ pockets; Asn142-Gly146 in the oxyanion hole loop, with the largest displacement observed in Gly143; His163 and Met165 whose sidechains form parts of the S1 and S2 pockets respectively; and Ala191-Gln192 which lie behind the substrate P5-P4 residues in the substrate complex.

**Figure 3:**
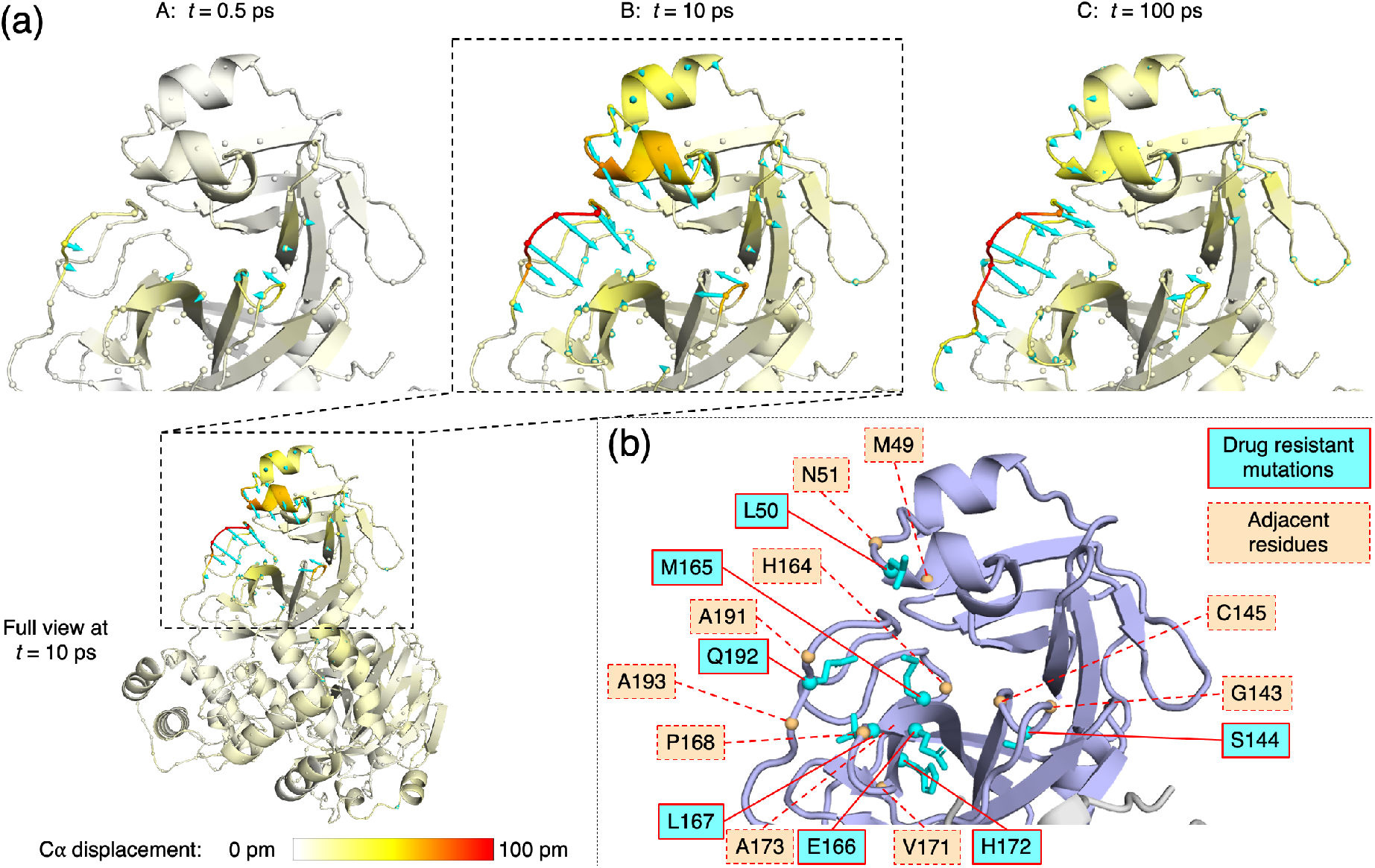
D-NEMD analysis of displacement vectors identifies sites associated with nirmatrelvir resistance. (a) Views of the M^pro^-s05 complex prior to MD simulations, with a focus on the substrate binding site, where significant D-NEMD responses, measured by averaging Cα displacement vectors between the equilibrium and nonequilibrium trajectories, are observed. Displacement magnitudes are represented on a white-yellow-red scale (figures created using PyMOL).^45^ Vectors with length ≥15 pm are displayed as cyan arrows with a scale-up factor of 5.^51^ (b) M^pro^ residues for which mutations associated with nirmatrelvir resistance have been reported,^52-54^ and adjacent residues in protein sequence.

**Table 1:**
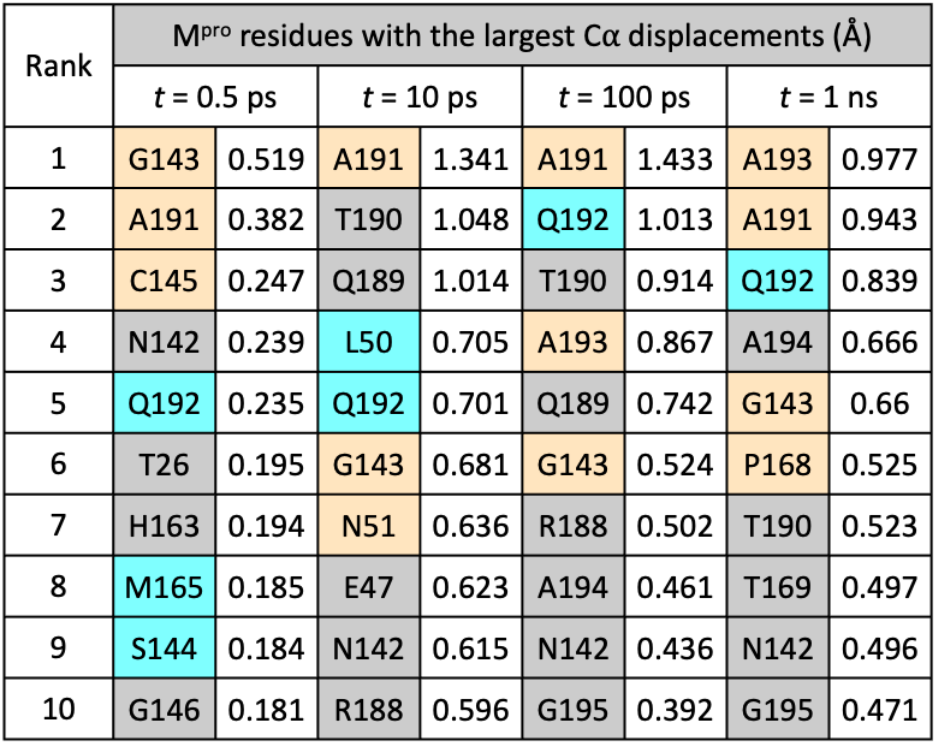
The most perturbed M^pro^ residues in terms of Cα displacements following substrate removal. Residues corresponding to and adjacent to mutation sites are coloured cyan and orange, respectively (see **Figure 3b**).

At *t* = 10 ps (**Figure 3aB; Table 1**), additional displacements are observed in the Ser46-Leu50 and Tyr54-Arg60 helices and their connecting loop above the active site in chain A. Displacements in residues surrounding Ala191 and Gly143 increase in magnitude, with the Arg188-Gln192 loop moving outwards into the substrate S5 and S4 pockets, and the Asn142-Gly143 loop closing inwards into S1. These motions collectively reflect a closure of the M^pro^ substrate binding site upon substrate removal.

From *t* = 10 ps to 100 ps (**Figure 3aC**), a reduction in the vector response is observed in certain regions, such as the Ser46-Leu50 helix, suggesting that there is no longer a consistent directional difference in the motions between the equilibrium and nonequilibrium trajectories. By contrast, the displacements in the Gly143 oxyanion hole loop and in the Val186-Gly195 loop persist. In the latter, the response is initially centred at Ala191, but gradually propagates downstream to residues as far as Gly195 (**Figures 3aC, S4.6; Table 1**). The loop in chain A remains the most perturbed region in the dimer at *t* = 5 ns (**Figures S4.5-6**), with Ala193 showing the largest displacement of 0.91 Å. These observations likely reflect long-lived structural alterations to accommodate the substrate. The conformational adaptation of the Val186-Gly195 loop observed here echoes structural studies of M^pro^ complexed with substrate peptides: the loop has crystallographically been found to be conformationally variable, which enables the binding of diverse substrate sequences.^12^ Overall, the D-NEMD approach provides a statistically meaningful prediction of how each M^pro^ region dynamically adapts to bind its substrate. The collective and coherent motion is not limited to the active site but extends well beyond.

### Conclusions

Selection pressure caused by treatments that target M^pro^, such as nirmatrelvir, drives the emergences of drug resistant mutations in SARS-CoV-2 M^pro^.^24, 34, 54, 55^ Such mutations would be expected to weaken the binding affinity of nirmatrelvir, but still allow M^pro^ to perform its biological function by binding, recognising, and hydrolysing its substrate sequences in the viral polyproteins. The effect of such mutations on dynamics is of particular interest. Indeed, M^pro^ mutations that confer high levels of resistance to nirmatrelvir have been reported, including the combined L50F+E166A+L167F substitutions,^52^ L50F+E166V substitutions,^53^ and naturally observed variations including S144M/F/A/G/Y, M165T, E166G, H172Q/F, and Q192T/S/L/A/I/P/H/V/W/C/F (**Figure 3b**).^54^ Some of these mutated residues, such as Glu166 and Met165, are in direct contact with the substrate P3/P1 and P2 residues respectively, and these interactions are exploited by nirmatrelvir.^9^ The effect of mutations that are more distant from the active site, such as those involving Leu50 and Gln192, on substrate and inhibitor binding, however, is less clear. The D-NEMD results here indicate that these residues are involved in allosteric response to active site binding.

Our D-NEMD results (**Figure 3a; Table 1**) highlight the dynamic behaviour of the oxyanion hole loop around Gly143. The dynamics of this loop will probably be affected by the observed mutation of Ser144, due to alterations in size, polarity, and HB interactions. Further from the catalytic site, the simulations identify Leu50 and its associated helix (Ser46-Leu50), as well as Gln192 and the loop of which it forms part, as being affected by the loss of substrate. While the exact influence of these mutations on binding affinities and reactivities of substrates or inhibitors remains to be investigated, this study has showcased the ability of the D-NEMD approach to provide valuable insights into time-ordered protein structural changes and pinpoint positions of drug-resistant mutations. The evaluation of the perturbation-induced responses also enables the identification, and potentially prediction, of allosteric sites that play major roles in regulating enzyme activity.

The D-NEMD approach is emerging as a useful tool to investigate signal propagation, identify communication networks, and characterise allosteric effects.^38^ As demonstrated by this study, D-NEMD simulations will be useful in predicting allosteric sites and mutation sites relevant to drug resistance.

## Methods

### Equilibrium MD

MD simulations were performed with GROMACS (v 2019.2)^56^ using the AMBER99SB-ILDN forcefield.^57^ Dimeric M^pro^ (PDB 6YB7;^58^ 1.25 Å resolution; chains A and B) in noncovalent complex with the s05 peptide was constructed and prepared as described previously, including the setup of histidine residues (**Table S5.1**) and the neutral catalytic dyad.^20^ The complex was placed in a rhombic dodecahedral box with at least 1.0 nm separation from box edges, solvated with TIP3P water,^59^ neutralised with sodium ions (86,304 atoms in solvated system), and minimised until the maximum force was below 1000 kJ mol^-1^ nm^-1^. From the minimised system, five replicas were initiated by assigning random velocities at 298.15 K. Each replica was subjected to 200 ps (1 fs step) NVT and 200 ps (1 fs step) NPT equilibration at 298.15 K and 1.0 bar, before a production run of 200 ns (2 fs step), during which coordinates were saved every 100 ps and velocities saved every 1 ns. Temperature coupling to the protein and non-protein at 298.15 K was achieved using a velocity-rescaling thermostat with a stochastic term, with a time constant of 0.1 ps.^60^ Pressure was maintained at 1.0 bar with a Parrinello-Rahman barostat with a time constant of 2 ps.^61, 62^ Long-range electrostatics were treated using smooth Particle-mesh Ewald with a 1 nm cut-off.^63, 64^ For van der Waals interactions, a 1 nm cut-off was used. For these 5 × 200 ns equilibrium simulations, analysis was performed using GROMACS tools (v 2019.2)^56^ and DSSP (v 2.0.4).^65, 66^ Hydrogen bonds were defined based on a combined distance (*d*_D-A_ ≤ 3.5 Å) and angle (∠(H-D-A) ≤ 30°) criteria.

### D-NEMD

From each of the five equilibrium trajectories, coordinates and velocities were extracted every 5 ns from *t* = 50 ns up to 195 ns, providing a total of 5 × 30 = 150 configurations for D-NEMD initiation. For the nonequilibrium simulations, the s05 peptide was instantaneously removed. No new molecules were added in its place. The nonequilibrium simulations were then started from these conformations. To capture events happening over different timescales, four length scales were chosen for the restarted MD simulations (2 fs step) with different saving frequencies of the protein coordinates: (i) over 2 ps saving every 0.05 ps; (ii) over 200 ps saving every 0.5 ps; (iii) over 1 ns saving every 10 ps; (iv) over 5 ns saving every 100 ps (only conducted for the nonequilibrium propagation, as the resulting time points coincided with the equilibrium trajectories).

### D-NEMD analysis

At equivalent delays after perturbation, the M^pro^ structure from the nonequilibrium propagation was fitted onto the corresponding equilibrium structure based on the 612 M^pro^ Cα atoms, before calculation of the scalar deviation and equilibrium-to-nonequilibrium displacement vector of every Cα atom. The response of each Cα atom was determined using the Kubo-Onsager approach, comparing the equilibrium and nonequilibrium simulations at equivalent points in time.^37, 38^ For each time, the Cα atom scalar deviation and displacement vector were averaged over the 150 replicas. According to the Kubo-Onsager relation, the averages yield estimates of the macroscopic time-dependent response of the system. The statistical significance was assessed by the standard error of the mean (SEM; *N* = 150). Average deviations and displacements were visualised using the M^pro^-s05 complex structure following energy minimisation and prior to any MD simulations, with the aid of PyMOL (v 2.3.0)^45^ and the modevectors.py script.^51^

## ASSOCIATED CONTENT

Supporting Information Available: System setup, detailed analysis of equilibrium and nonequilibrium MD simulations.

## Data Availability

All simulation data (including input, structure, and trajectory files) are openly available on GitHub (https://github.com/duartegroup/Mpro-Substrate_D-NEMD).

## Acknowledgements

HTHC thanks the Clarendon Fund, New College Oxford, and the EPSRC Centre for Doctoral Training in Synthesis for Biology and Medicine (EP/L015838/1) for a studentship, generously supported by AstraZeneca, Diamond Light Source, Defence Science and Technology Laboratory, Evotec, GlaxoSmithKline, Janssen, Novartis, Pfizer, Syngenta, Takeda, UCB and Vertex. AJM and ASFO thank EPSRC (grant number EP/M022609/1), BBSRC (grant number BB/R016445/1), and ERC (Advanced Grant PREDACTED https://cordis.europa.eu/project/id/101021207) for support (funding from the European Research Council (ERC) under the European Union’s Horizon 2020 research and innovation programme, Grant agreement No. 101021207). This project made use of time on HPC granted via the UK High-End Computing Consortium for Biomolecular Simulation, HECBioSim (http://hecbiosim.ac.uk), supported by EPSRC (grant no. EP/R029407/1).

---

**For Table of Contents Only**

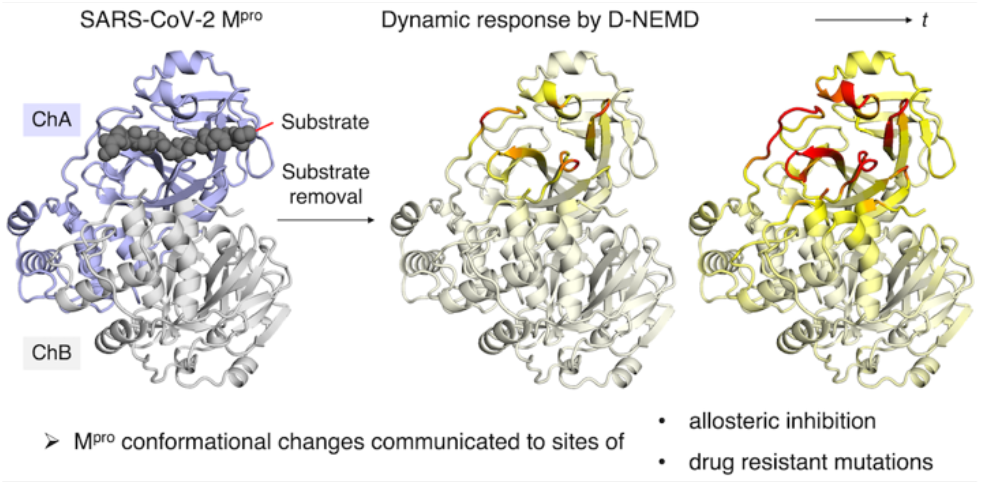

## References

1. Rut, W.; Groborz, K.; Zhang, L.; Sun, X.; Zmudzinski, M.; Pawlik, B.; Wang, X.; Jochmans, D.; Neyts, J.; Młynarski, W.; Hilgenfeld, R.; Drag, M. SARS-CoV-2 Mpro inhibitors and activity-based probes for patient-sample imaging. Nat. Chem. Biol. 2021, 17 (2), 222–228.

2. Wu, F.; Zhao, S.; Yu, B.; Chen, Y.-M.; Wang, W.; Song, Z.-G.; Hu, Y.; Tao, Z.-W.; Tian, J.-H.; Pei, Y.-Y.; Yuan, M.-L.; Zhang, Y.-L.; Dai, F.-H.; Liu, Y.; Wang, Q.-M.; Zheng, J.-J.; Xu, L.; Holmes, E. C.; Zhang, Y.-Z. A new coronavirus associated with human respiratory disease in China. Nature 2020, 579 (7798), 265–269.

3. Koudelka, T.; Boger, J.; Henkel, A.; Schönherr, R.; Krantz, S.; Fuchs, S.; Rodríguez, E.; Redecke, L.; Tholey, A. N-Terminomics for the Identification of In Vitro Substrates and Cleavage Site Specificity of the SARS-CoV-2 Main Protease. Proteomics 2021, 21 (2), 2000246.

4. MacDonald, E. A.; Frey, G.; Namchuk, M. N.; Harrison, S. C.; Hinshaw, S. M.; Windsor, I. W. Recognition of Divergent Viral Substrates by the SARS-CoV-2 Main Protease. ACS Infect. Dis. 2021, 7 (9), 2591–2595.

5. Jin, Z.; Du, X.; Xu, Y.; Deng, Y.; Liu, M.; Zhao, Y.; Zhang, B.; Li, X.; Zhang, L.; Peng, C.; Duan, Y.; Yu, J.; Wang, L.; Yang, K.; Liu, F.; Jiang, R.; Yang, X.; You, T.; Liu, X.; Yang, X.; Bai, F.; Liu, H.; Liu, X.; Guddat, L. W.; Xu, W.; Xiao, G.; Qin, C.; Shi, Z.; Jiang, H.; Rao, Z.; Yang, H. Structure of Mpro from SARS-CoV-2 and discovery of its inhibitors. Nature 2020, 582 (7811), 289–293.

6. Zhang, L.; Lin, D.; Sun, X.; Curth, U.; Drosten, C.; Sauerhering, L.; Becker, S.; Rox, K.; Hilgenfeld, R. Crystal structure of SARS-CoV-2 main protease provides a basis for design of improved α-ketoamide inhibitors. Science 2020, 368 (6489), 409–412.

7. Cho, E.; Rosa, M.; Anjum, R.; Mehmood, S.; Soban, M.; Mujtaba, M.; Bux, K.; Moin, S. T.; Tanweer, M.; Dantu, S.; Pandini, A.; Yin, J.; Ma, H.; Ramanathan, A.; Islam, B.; Mey, A. S. J. S.; Bhowmik, D.; Haider, S. Dynamic Profiling of β-Coronavirus 3CL Mpro Protease Ligand-Binding Sites. J. Chem. Inf. Model. 2021, 61 (6), 3058–3073.

8. Lv, Z.; Cano, K. E.; Jia, L.; Drag, M.; Huang, T. T.; Olsen, S. K. Targeting SARS-CoV-2 Proteases for COVID-19 Antiviral Development. Front. Chem. 2022, 9, 819165.

9. Owen, D. R.; Allerton, C. M. N.; Anderson, A. S.; Aschenbrenner, L.; Avery, M.; Berritt, S.; Boras, B.; Cardin, R. D.; Carlo, A.; Coffman, K. J.; Dantonio, A.; Di, L.; Eng, H.; Ferre, R.; Gajiwala, K. S.; Gibson, S. A.; Greasley, S. E.; Hurst, B. L.; Kadar, E. P.; Kalgutkar, A. S.; Lee, J. C.; Lee, J.; Liu, W.; Mason, S. W.; Noell, S.; Novak, J. J.; Obach, R. S.; Ogilvie, K.; Patel, N. C.; Pettersson, M.; Rai, D. K.; Reese, M. R.; Sammons, M. F.; Sathish, J. G.; Singh, R. S. P.; Steppan, C. M.; Stewart, A. E.; Tuttle, J. B.; Updyke, L.; Verhoest, P. R.; Wei, L.; Yang, Q.; Zhu, Y. An oral SARS-CoV-2 Mpro inhibitor clinical candidate for the treatment of COVID-19. Science 2021, 374 (6575), 1586–1593.

10. US Food & Drug Administration, Coronavirus (COVID-19) Update: FDA Authorizes First Oral Antiviral for Treatment of COVID-19. US FDA: December 22, 2021.

11. Unoh, Y.; Uehara, S.; Nakahara, K.; Nobori, H.; Yamatsu, Y.; Yamamoto, S.; Maruyama, Y.; Taoda, Y.; Kasamatsu, K.; Suto, T.; Kouki, K.; Nakahashi, A.; Kawashima, S.; Sanaki, T.; Toba, S.; Uemura, K.; Mizutare, T.; Ando, S.; Sasaki, M.; Orba, Y.; Sawa, H.; Sato, A.; Sato, T.; Kato, T.; Tachibana, Y. Discovery of S-217622, a Noncovalent Oral SARS-CoV-2 3CL Protease Inhibitor Clinical Candidate for Treating COVID-19. J. Med. Chem. 2022, 65 (9), 6499–6512.

12. Shaqra, A. M.; Zvornicanin, S. N.; Huang, Q. Y. J.; Lockbaum, G. J.; Knapp, M.; Tandeske, L.; Bakan, D. T.; Flynn, J.; Bolon, D. N. A.; Moquin, S.; Dovala, D.; Kurt Yilmaz, N.; Schiffer, C. A. Defining the substrate envelope of SARS-CoV-2 main protease to predict and avoid drug resistance. Nat. Commun. 2022, 13 (1), 3556.

13. Lee, J.; Worrall, L. J.; Vuckovic, M.; Rosell, F. I.; Gentile, F.; Ton, A.-T.; Caveney, N. A.; Ban, F.; Cherkasov, A.; Paetzel, M.; Strynadka, N. C. J. Crystallographic structure of wild-type SARS-CoV-2 main protease acyl-enzyme intermediate with physiological C-terminal autoprocessing site. Nat. Commun. 2020, 11 (1), 5877.

14. Kneller, D. W.; Zhang, Q.; Coates, L.; Louis, J. M.; Kovalevsky, A. Michaelis-like complex of SARS-CoV-2 main protease visualized by room-temperature X-ray crystallography. IUCrJ 2021, 8 (6), 973–979.

15. Suárez, D.; Díaz, N. SARS-CoV-2 Main Protease: A Molecular Dynamics Study. J. Chem. Inf. Model. 2020, 60 (12), 5815–5831.

16. Díaz, N.; Suárez, D. Influence of charge configuration on substrate binding to SARS-CoV-2 main protease. Chem. Commun. 2021, 57 (43), 5314–5317.

17. Świderek, K.; Moliner, V. Revealing the molecular mechanisms of proteolysis of SARS-CoV-2 Mpro by QM/MM computational methods. Chem. Sci. 2020, 11 (39), 10626–10630.

18. Ramos-Guzmán, C. A.; Ruiz-Pernía, J. J.; Tuñón, I. Unraveling the SARS-CoV-2 Main Protease Mechanism Using Multiscale Methods. ACS Catal. 2020, 10 (21), 12544–12554.

19. Khrenova, M. G.; Tsirelson, V. G.; Nemukhin, A. V. Dynamical properties of enzyme–substrate complexes disclose substrate specificity of the SARS-CoV-2 main protease as characterized by the electron density descriptors. Phys. Chem. Chem. Phys. 2020, 22 (34), 19069–19079.

20. Chan, H. T. H.; Moesser, M. A.; Walters, R. K.; Malla, T. R.; Twidale, R. M.; John, T.; Deeks, H. M.; Johnston-Wood, T.; Mikhailov, V.; Sessions, R. B.; Dawson, W.; Salah, E.; Lukacik, P.; Strain-Damerell, C.; Owen, C. D.; Nakajima, T.; Świderek, K.; Lodola, A.; Moliner, V.; Glowacki, D. R.; Spencer, J.; Walsh, M. A.; Schofield, C. J.; Genovese, L.; Shoemark, D. K.; Mulholland, A. J.; Duarte, F.; Morris, G. M. Discovery of SARS-CoV-2 Mpro peptide inhibitors from modelling substrate and ligand binding. Chem. Sci. 2021, 12 (41), 13686–13703.

21. Sztain, T.; Amaro, R.; McCammon, J. A. Elucidation of Cryptic and Allosteric Pockets within the SARS-CoV-2 Main Protease. J. Chem. Inf. Model. 2021, 61 (7), 3495–3501.

22. Arafet, K.; Serrano-Aparicio, N.; Lodola, A.; Mulholland, A. J.; González, F. V.; Świderek, K.; Moliner, V. Mechanism of inhibition of SARS-CoV-2 Mpro by N3 peptidyl Michael acceptor explained by QM/MM simulations and design of new derivatives with tunable chemical reactivity. Chem. Sci. 2021, 12 (4), 1433–1444.

23. Martí, S.; Arafet, K.; Lodola, A.; Mulholland, A. J.; Świderek, K.; Moliner, V. Impact of Warhead Modulations on the Covalent Inhibition of SARS-CoV-2 Mpro Explored by QM/MM Simulations. ACS Catal. 2022, 12 (1), 698–708.

24. Ramos-Guzmán, C. A.; Andjelkovic, M.; Zinovjev, K.; Ruiz-Pernía, J. J.; Tuñón, I. The impact of SARS-CoV-2 3CL protease mutations on nirmatrelvir inhibitory efficiency. Computational insights into potential resistance mechanisms. Chem. Sci. 2023, 14 (10), 2686–2697.

25. El Khoury, L.; Jing, Z.; Cuzzolin, A.; Deplano, A.; Loco, D.; Sattarov, B.; Hédin, F.; Wendeborn, S.; Ho, C.; El Ahdab, D.; Jaffrelot Inizan, T.; Sturlese, M.; Sosic, A.; Volpiana, M.; Lugato, A.; Barone, M.; Gatto, B.; Macchia, M. L.; Bellanda, M.; Battistutta, R.; Salata, C.; Kondratov, I.; Iminov, R.; Khairulin, A.; Mykhalonok, Y.; Pochepko, A.; Chashka-Ratushnyi, V.; Kos, I.; Moro, S.; Montes, M.; Ren, P.; Ponder, J. W.; Lagardère, L.; Piquemal, J.-P.; Sabbadin, D. Computationally driven discovery of SARS-CoV-2 Mpro inhibitors: from design to experimental validation. Chem. Sci. 2022, 13 (13), 3674–3687.

26. Thakur, A.; Sharma, G.; Badavath, V. N.; Jayaprakash, V.; Merz, K. M., Jr.; Blum, G.; Acevedo, O. Primer for Designing Main Protease (Mpro) Inhibitors of SARS-CoV-2. J. Phys. Chem. Lett. 2022, 13 (25), 5776–5786.

27. Matthew, A. N.; Leidner, F.; Lockbaum, G. J.; Henes, M.; Zephyr, J.; Hou, S.; Rao, D. N.; Timm, J.; Rusere, L. N.; Ragland, D. A.; Paulsen, J. L.; Prachanronarong, K.; Soumana, D. I.; Nalivaika, E. A.; Kurt Yilmaz, N.; Ali, A.; Schiffer, C. A. Drug Design Strategies to Avoid Resistance in Direct-Acting Antivirals and Beyond. Chem. Rev. 2021, 121 (6), 3238–3270.

28. King, N. M.; Prabu-Jeyabalan, M.; Bandaranayake, R. M.; Nalam, M. N.; Nalivaika, E. A.; Özen, A.; Haliloğlu, T.; Yilmaz, N. K.; Schiffer, C. A. Extreme entropy-enthalpy compensation in a drug-resistant variant of HIV-1 protease. ACS Chem. Biol. 2012, 7 (9), 1536–46.

29. Callegari, D.; Ranaghan, K. E.; Woods, C. J.; Minari, R.; Tiseo, M.; Mor, M.; Mulholland, A. J.; Lodola, A. L718Q mutant EGFR escapes covalent inhibition by stabilizing a non-reactive conformation of the lung cancer drug osimertinib. Chem. Sci. 2018, 9 (10), 2740–2749.

30. Galdadas, I.; Qu, S.; Oliveira, A. S. F.; Olehnovics, E.; Mack, A. R.; Mojica, M. F.; Agarwal, P. K.; Tooke, C. L.; Gervasio, F. L.; Spencer, J.; Bonomo, R. A.; Mulholland, A. J.; Haider, S. Allosteric communication in class A β-lactamases occurs via cooperative coupling of loop dynamics. eLife 2021, 10, e66567.

31. Henes, M.; Lockbaum, G. J.; Kosovrasti, K.; Leidner, F.; Nachum, G. S.; Nalivaika, E. A.; Lee, S. K.; Spielvogel, E.; Zhou, S.; Swanstrom, R.; Bolon, D. N. A.; Kurt Yilmaz, N.; Schiffer, C. A. Picomolar to Micromolar: Elucidating the Role of Distal Mutations in HIV-1 Protease in Conferring Drug Resistance. ACS Chem. Biol. 2019, 14 (11), 2441–2452.

32. Moghadasi, S. A.; Esler, M. A.; Otsuka, Y.; Becker, J. T.; Moraes, S. N.; Anderson, C. B.; Chamakuri, S.; Belica, C.; Wick, C.; Harki, D. A.; Young, D. W.; Scampavia, L.; Spicer, T. P.; Shi, K.; Aihara, H.; Brown, W. L.; Harris, R. S. Gain-of-Signal Assays for Probing Inhibition of SARS-CoV-2 Mpro/3CLpro in Living Cells. mBio 2022, 13 (3), e00784–22.

33. Telenti, A.; Hodcroft, E. B.; Robertson, D. L. The Evolution and Biology of SARS-CoV-2 Variants. Cold Spring Harb. Perspect. Med. 2022, 12 (5), a041390.

34. Flynn, J. M.; Samant, N.; Schneider-Nachum, G.; Barkan, D. T.; Yilmaz, N. K.; Schiffer, C. A.; Moquin, S. A.; Dovala, D.; Bolon, D. N. A. Comprehensive fitness landscape of SARS-CoV-2 Mpro reveals insights into viral resistance mechanisms. eLife 2022, 11, e77433.

35. Ciccotti, G.; Jacucci, G. Direct Computation of Dynamical Response by Molecular Dynamics: The Mobility of a Charged Lennard-Jones Particle. Phys. Rev. Lett. 1975, 35 (12), 789–792.

36. Ciccotti, G.; Jacucci, G.; McDonald, I. R. “Thought-experiments” by molecular dynamics. J. Stat. Phys. 1979, 21 (1), 1–22.

37. Ciccotti, G.; Ferrario, M. Non-equilibrium by molecular dynamics: a dynamical approach. Mol. Simul. 2016, 42 (16), 1385–1400.

38. Oliveira, A. S. F.; Ciccotti, G.; Haider, S.; Mulholland, A. J. Dynamical nonequilibrium molecular dynamics reveals the structural basis for allostery and signal propagation in biomolecular systems. Eur. Phys. J. B 2021, 94 (7), 144.

39. Oliveira, A. S. F.; Shoemark, D. K.; Campello, H. R.; Wonnacott, S.; Gallagher, T.; Sessions, R. B.; Mulholland, A. J. Identification of the Initial Steps in Signal Transduction in the α4β2 Nicotinic Receptor: Insights from Equilibrium and Nonequilibrium Simulations. Structure 2019, 27 (7), 1171–1183.e3.

40. Oliveira, A. S. F.; Edsall, C. J.; Woods, C. J.; Bates, P.; Nunez, G. V.; Wonnacott, S.; Bermudez, I.; Ciccotti, G.; Gallagher, T.; Sessions, R. B.; Mulholland, A. J. A General Mechanism for Signal Propagation in the Nicotinic Acetylcholine Receptor Family. J. Am. Chem. Soc. 2019, 141 (51), 19953–19958.

41. Dommer, A.; Casalino, L.; Kearns, F.; Rosenfeld, M.; Wauer, N.; Ahn, S.-H.; Russo, J.; Oliveira, S.; Morris, C.; Bogetti, A.; Trifan, A.; Brace, A.; Sztain, T.; Clyde, A.; Ma, H.; Chennubhotla, C.; Lee, H.; Turilli, M.; Khalid, S.; Tamayo-Mendoza, T.; Welborn, M.; Christensen, A.; Smith, D. G.; Qiao, Z.; Sirumalla, S. K.; O’Connor, M.; Manby, F.; Anandkumar, A.; Hardy, D.; Phillips, J.; Stern, A.; Romero, J.; Clark, D.; Dorrell, M.; Maiden, T.; Huang, L.; McCalpin, J.; Woods, C.; Gray, A.; Williams, M.; Barker, B.; Rajapaksha, H.; Pitts, R.; Gibbs, T.; Stone, J.; Zuckerman, D. M.; Mulholland, A. J.; Miller, T.; Jha, S.; Ramanathan, A.; Chong, L.; Amaro, R. E. #COVIDisAirborne: AI-enabled multiscale computational microscopy of delta SARS-CoV-2 in a respiratory aerosol. Int. J. High Perform. Comput. Appl. 2022, 0 (0), 10943420221128233.

42. Oliveira, A. S. F.; Shoemark, D. K.; Avila Ibarra, A.; Davidson, A. D.; Berger, I.; Schaffitzel, C.; Mulholland, A. J. The fatty acid site is coupled to functional motifs in the SARS-CoV-2 spike protein and modulates spike allosteric behaviour. Comput. Struct. Biotechnol. J. 2022, 20, 139–147.

43. Oliveira, A. S. F.; Shoemark, D. K.; Davidson, A. D.; Berger, I.; Schaffitzel, C.; Mulholland, A. J. SARS-CoV-2 spike variants differ in their allosteric response to linoleic acid. J. Mol. Cell. Biol. 2023, mjad021.

44. Gupta, K.; Toelzer, C.; Williamson, M. K.; Shoemark, D. K.; Oliveira, A. S. F.; Matthews, D. A.; Almuqrin, A.; Staufer, O.; Yadav, S. K. N.; Borucu, U.; Garzoni, F.; Fitzgerald, D.; Spatz, J.; Mulholland, A. J.; Davidson, A. D.; Schaffitzel, C.; Berger, I. Structural insights in cell-type specific evolution of intra-host diversity by SARS-CoV-2. Nat. Commun. 2022, 13 (1), 222.

45. Schrödinger LLC. The PyMOL Molecular Graphics System, Version 2.3.0.

46. Shi, J.; Han, N.; Lim, L.; Lua, S.; Sivaraman, J.; Wang, L.; Mu, Y.; Song, J. Dynamically-Driven Inactivation of the Catalytic Machinery of the SARS 3C-Like Protease by the N214A Mutation on the Extra Domain. PLOS Comput. Biol. 2011, 7 (2), e1001084.

47. Kneller, D. W.; Phillips, G.; O’Neill, H. M.; Jedrzejczak, R.; Stols, L.; Langan, P.; Joachimiak, A.; Coates, L.; Kovalevsky, A. Structural plasticity of SARS-CoV-2 3CL Mpro active site cavity revealed by room temperature X-ray crystallography. Nat. Commun. 2020, 11 (1), 3202.

48. Günther, S.; Reinke, P. Y. A.; Fernández-García, Y.; Lieske, J.; Lane, T. J.; Ginn, H. M.; Koua, F. H. M.; Ehrt, C.; Ewert, W.; Oberthuer, D.; Yefanov, O.; Meier, S.; Lorenzen, K.; Krichel, B.; Kopicki, J.-D.; Gelisio, L.; Brehm, W.; Dunkel, I.; Seychell, B.; Gieseler, H.; Norton-Baker, B.; Escudero-Pérez, B.; Domaracky, M.; Saouane, S.; Tolstikova, A.; White, T. A.; Hänle, A.; Groessler, M.; Fleckenstein, H.; Trost, F.; Galchenkova, M.; Gevorkov, Y.; Li, C.; Awel, S.; Peck, A.; Barthelmess, M.; Schlünzen, F.; Lourdu Xavier, P.; Werner, N.; Andaleeb, H.; Ullah, N.; Falke, S.; Srinivasan, V.; França, B. A.; Schwinzer, M.; Brognaro, H.; Rogers, C.; Melo, D.; Zaitseva-Doyle, J. J.; Knoska, J.; Peña-Murillo, G. E.; Mashhour, A. R.; Hennicke, V.; Fischer, P.; Hakanpää, J.; Meyer, J.; Gribbon, P.; Ellinger, B.; Kuzikov, M.; Wolf, M.; Beccari, A. R.; Bourenkov, G.; von Stetten, D.; Pompidor, G.; Bento, I.; Panneerselvam, S.; Karpics, I.; Schneider, T. R.; Garcia-Alai, M. M.; Niebling, S.; Günther, C.; Schmidt, C.; Schubert, R.; Han, H.; Boger, J.; Monteiro, D. C. F.; Zhang, L.; Sun, X.; Pletzer-Zelgert, J.; Wollenhaupt, J.; Feiler, C. G.; Weiss, M. S.; Schulz, E.-C.; Mehrabi, P.; Karničar, K.; Usenik, A.; Loboda, J.; Tidow, H.; Chari, A.; Hilgenfeld, R.; Uetrecht, C.; Cox, R.; Zaliani, A.; Beck, T.; Rarey, M.; Günther, S.; Turk, D.; Hinrichs, W.; Chapman, H. N.; Pearson, A. R.; Betzel, C.; Meents, A. X-ray screening identifies active site and allosteric inhibitors of SARS-CoV-2 main protease. Science 2021, 372 (6542), 642–646.

49. Alzyoud, L.; Ghattas, M. A.; Atatreh, N. Allosteric Binding Sites of the SARS-CoV-2 Main Protease: Potential Targets for Broad-Spectrum Anti-Coronavirus Agents. Drug. Des. Devel. Ther. 2022, 16, 2463–2478.

50. Abreu, B.; Lopes, E. F.; Oliveira, A. S. F.; Soares, C. M. F508del disturbs the dynamics of the nucleotide binding domains of CFTR before and after ATP hydrolysis. Proteins 2020, 88 (1), 113–126.

51. Law, S. M. modevectors.py, https://pymolwiki.org/index.php/Modevectors, 2012 (accessed 2022-06-30).

52. Jochmans, D.; Liu, C.; Donckers, K.; Stoycheva, A.; Boland, S.; Stevens, S. K.; Vita, C. D.; Vanmechelen, B.; Maes, P.; Trüeb, B.; Ebert, N.; Thiel, V.; Jonghe, S. D.; Vangeel, L.; Bardiot, D.; Jekle, A.; Blatt, L. M.; Beigelman, L.; Symons, J. A.; Raboisson, P.; Chaltin, P.; Marchand, A.; Neyts, J.; Deval, J.; Vandyck, K. The Substitutions L50F, E166A, and L167F in SARS-CoV-2 3CLpro Are Selected by a Protease Inhibitor In Vitro and Confer Resistance To Nirmatrelvir. mBio 2023, 14 (1), e02815–22.

53. Zhou, Y.; Gammeltoft, K. A.; Ryberg, L. A.; Pham, L. V.; Tjørnelund, H. D.; Binderup, A.; Duarte Hernandez, C. R.; Fernandez-Antunez, C.; Offersgaard, A.; Fahnøe, U.; Peters, G. H. J.; Ramirez, S.; Bukh, J.; Gottwein, J. M. Nirmatrelvir-resistant SARS-CoV-2 variants with high fitness in an infectious cell culture system. Sci. Adv. 2022, 8 (51), eadd7197.

54. Hu, Y.; Lewandowski, E. M.; Tan, H.; Zhang, X.; Morgan, R. T.; Zhang, X.; Jacobs, L. M. C.; Butler, S. G.; Gongora, M. V.; Choy, J.; Deng, X.; Chen, Y.; Wang, J. Naturally occurring mutations of SARS-CoV-2 main protease confer drug resistance to nirmatrelvir. bioRxiv 2022, 2022.06.28.497978.

55. Moghadasi, S. A.; Heilmann, E.; Khalil, A. M.; Nnabuife, C.; Kearns, F. L.; Ye, C.; Moraes, S. N.; Costacurta, F.; Esler, M. A.; Aihara, H.; von Laer, D.; Martinez-Sobrido, L.; Palzkill, T.; Amaro, R. E.; Harris, R. S. Transmissible SARS-CoV-2 variants with resistance to clinical protease inhibitors. Sci. Adv. 2023, 9 (13), eade8778.

56. Abraham, M. J.; Murtola, T.; Schulz, R.; Páll, S.; Smith, J. C.; Hess, B.; Lindahl, E. GROMACS: High performance molecular simulations through multi-level parallelism from laptops to supercomputers. SoftwareX 2015, 1-2, 19–25.

57. Lindorff-Larsen, K.; Piana, S.; Palmo, K.; Maragakis, P.; Klepeis, J. L.; Dror, R. O.; Shaw, D. E. Improved side-chain torsion potentials for the Amber ff99SB protein force field. Proteins 2010, 78 (8), 1950–1958.

58. Owen, C. D.; Lukacik, P.; Strain-Damerell, C. M.; Douangamath, A.; Powell, A. J.; Fearon, D.; Brandao-Neto, J.; Crawshaw, A. D.; Aragao, D.; Williams, M.; Flaig, R.; Hall, D.; McAauley, K.; Stuart, D. I.; von Delft, F.; Walsh, M. A. COVID-19 main protease with unliganded active site. To be published 2020, PDB 6YB7, doi: 10.2210/pdb6yb7/pdb.

59. Jorgensen, W. L.; Chandrasekhar, J.; Madura, J. D.; Impey, R. W.; Klein, M. L. Comparison of simple potential functions for simulating liquid water. J. Chem. Phys. 1983, 79 (2), 926–935.

60. Bussi, G.; Donadio, D.; Parrinello, M. Canonical sampling through velocity rescaling. J. Chem. Phys. 2007, 126 (1), 014101.

61. Parrinello, M.; Rahman, A. Polymorphic transitions in single crystals: A new molecular dynamics method. J. Appl. Phys. 1981, 52 (12), 7182–7190.

62. Nosé, S.; Klein, M. L. Constant pressure molecular dynamics for molecular systems. Mol. Phys. 1983, 50 (5), 1055–1076.

63. Darden, T.; York, D.; Pedersen, L. Particle mesh Ewald: An N⋅log(N) method for Ewald sums in large systems. J. Chem. Phys. 1993, 98 (12), 10089–10092.

64. Essmann, U.; Perera, L.; Berkowitz, M. L.; Darden, T.; Lee, H.; Pedersen, L. G. A smooth particle mesh Ewald method. J. Chem. Phys. 1995, 103 (19), 8577–8593.

65. Touw, W. G.; Baakman, C.; Black, J.; te Beek, T. A. H.; Krieger, E.; Joosten, R. P.; Vriend, G. A series of PDB-related databanks for everyday needs. Nucleic Acids Res. 2015, 43 (Database issue), D364–D368.

66. Kabsch, W.; Sander, C. Dictionary of protein secondary structure: Pattern recognition of hydrogen-bonded and geometrical features. Biopolymers 1983, 22 (12), 2577–2637.

