## Supplementary material for "Dynamical nonequilibrium molecular dynamics simulations identify allosteric sites and positions associated with drug resistance in the SARS-CoV-2 main protease": SI

### Section S1: Equilibrium simulations of the M$^{pro}$-substrate complex

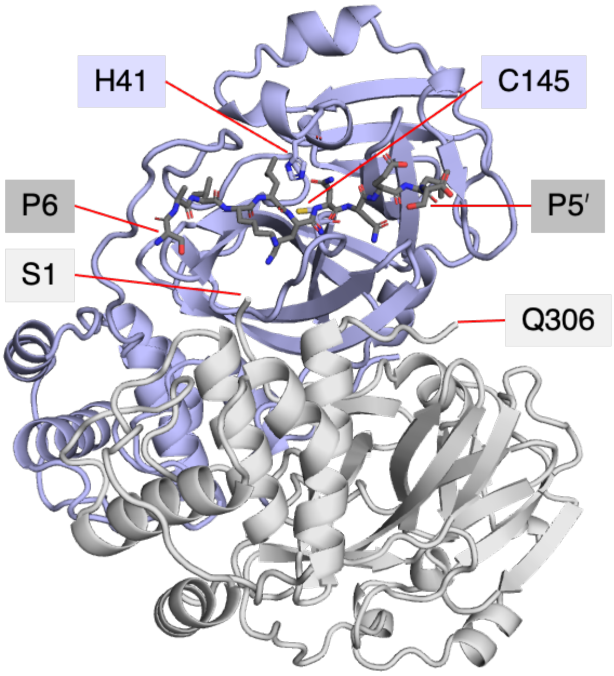


**Figure S1.1**: View of the starting non-covalent complex of dimeric M$^{pro}$ (chains A and B shown as blue and grey cartoons respectively, with the active site His41 and Cys145 labelled and shown as sticks; PDB 6YB7)[1] and the comparatively modelled s05 peptide (shown as dark grey sticks) representing the nsp8/9 native cleavage sequence.[2] Hydrogens are omitted for clarity. Chains A and B are subsequently referred to as ChA and ChB respectively. The N- and C-terminal residues of ChB are labelled.

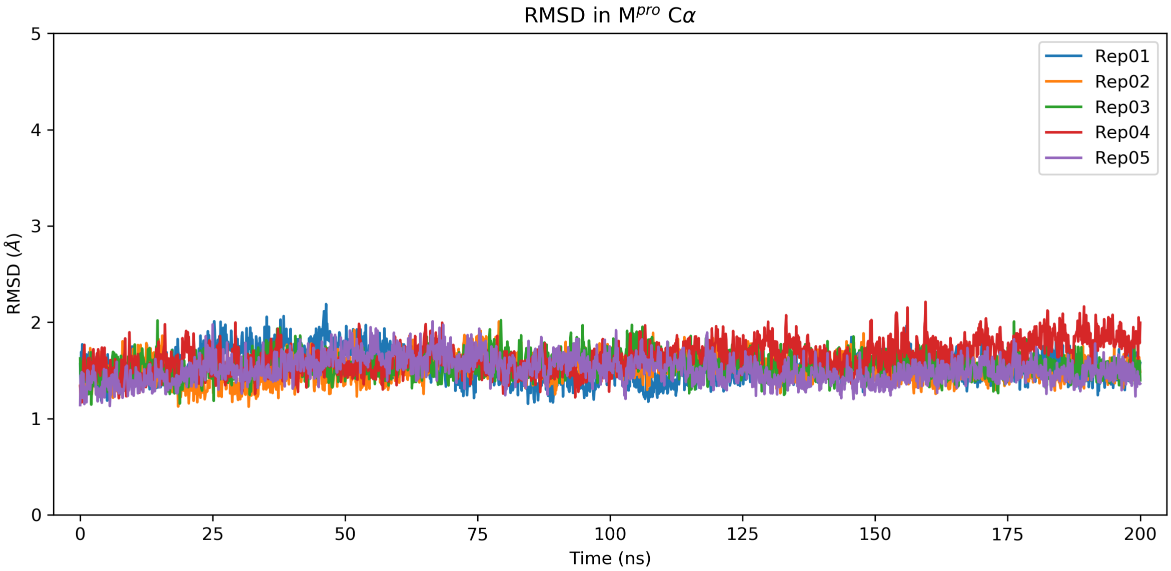


**Figure S1.2**: Time evolution of M$^{pro}$ Cα RMSD in each of the five 200 ns production MD simulations relative to the protein crystal structure. Each replica was subjected to 200 ps NVT equilibration, during which the protein and peptide heavy atoms were restrained, followed by 200 ps NPT equilibration where the restraints were released.

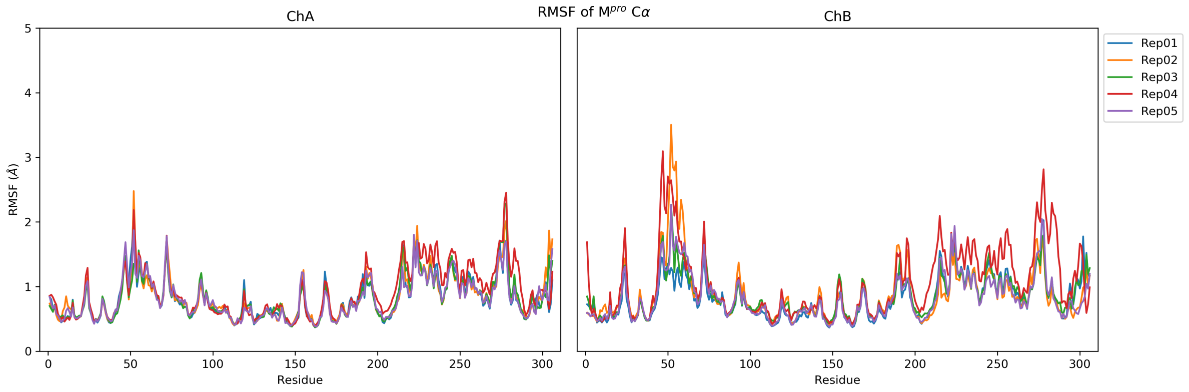


**Figure S1.3**: $M^{pro}$ Cα RMSF in each of the five 150 ns equilibrium MD simulations. The first 50 ns of each 200 ns production MD simulation was discarded as reflecting equilibration.

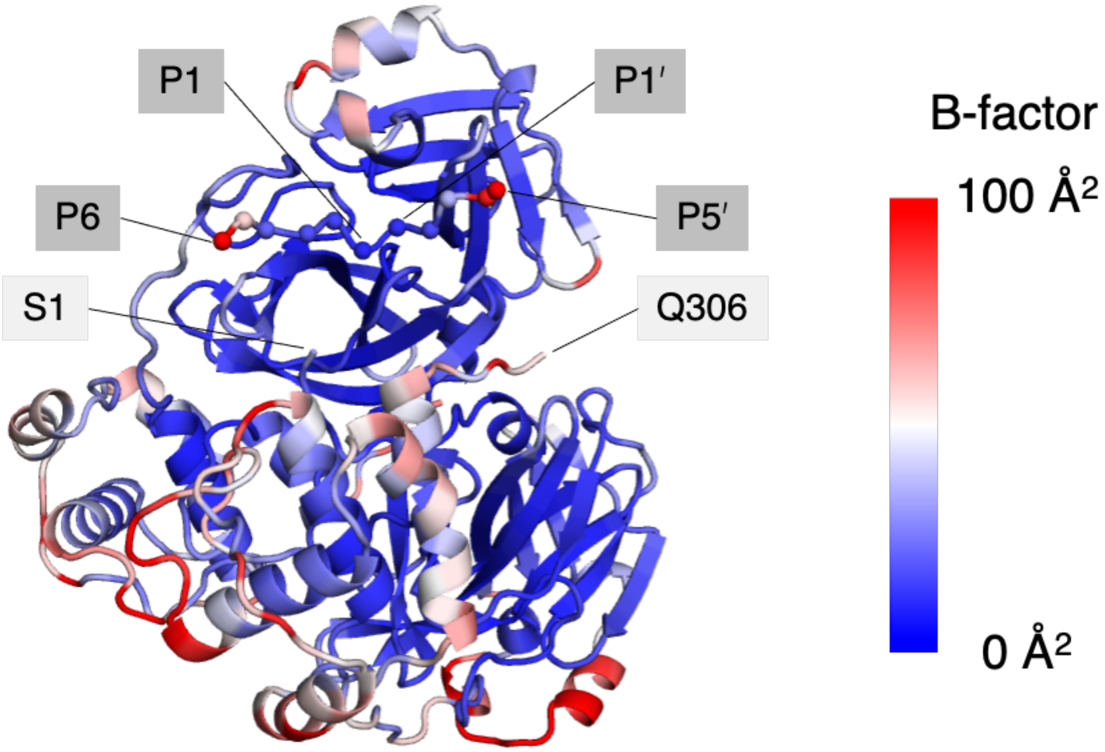


**Figure S1.4**: View of the global average structure of the $M^{pro}$-s05 complex over the 5 × 150 ns equilibrium MD simulations. The colouring is based on a 0-100 Å² blue-white-red scale of the MD-derived B-factors, with rigid regions in blue and flexible regions in red. The N- and C-terminal residues of ChB are labelled. The s05 peptide Cα atoms are shown as spheres, with the terminal (P6 and P5′) and the scissile amide (between P1 and P1′) positions labelled.

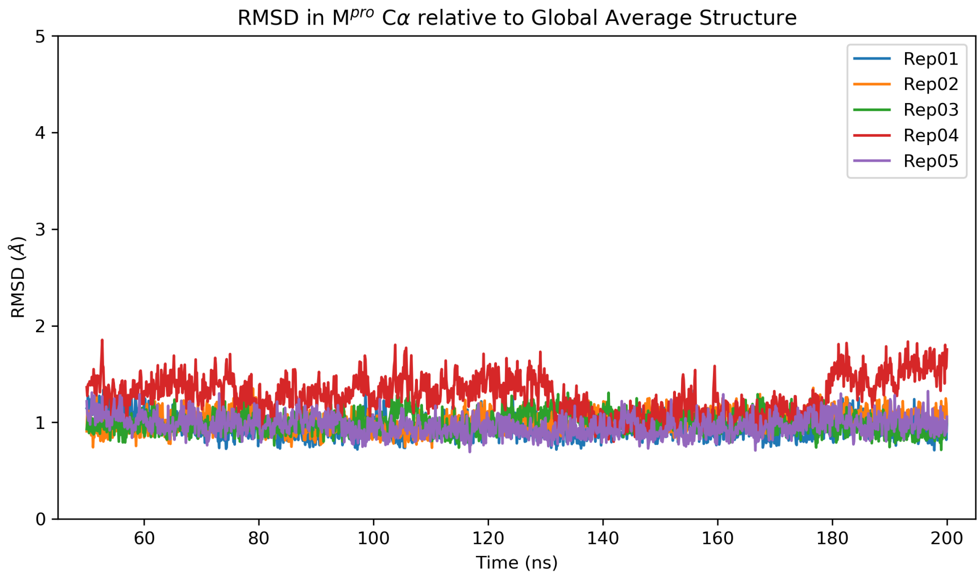


**Figure S1.5**: Time evolution of $M^{pro}$ Cα RMSD in each of the five 150 ns equilibrium MD simulations (starting from $t$ = 50 ns), relative to the equilibrium MD-derived global average structure of the complex.

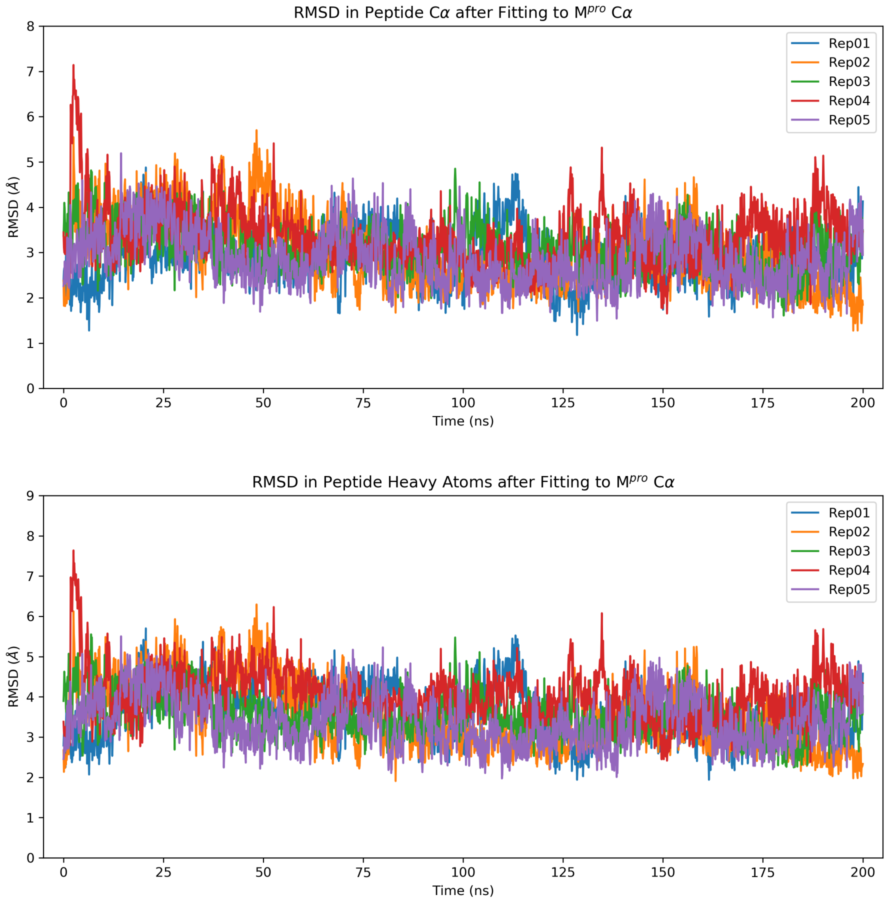


**Figure S1.6**: Time evolution of the s05 substrate peptide (top) Cα and (bottom) heavy atom RMSDs over the 5 × 200 ns production MD simulations relative to the starting conformation, with trajectories fitted based on $M^{pro}$ Cα atoms.

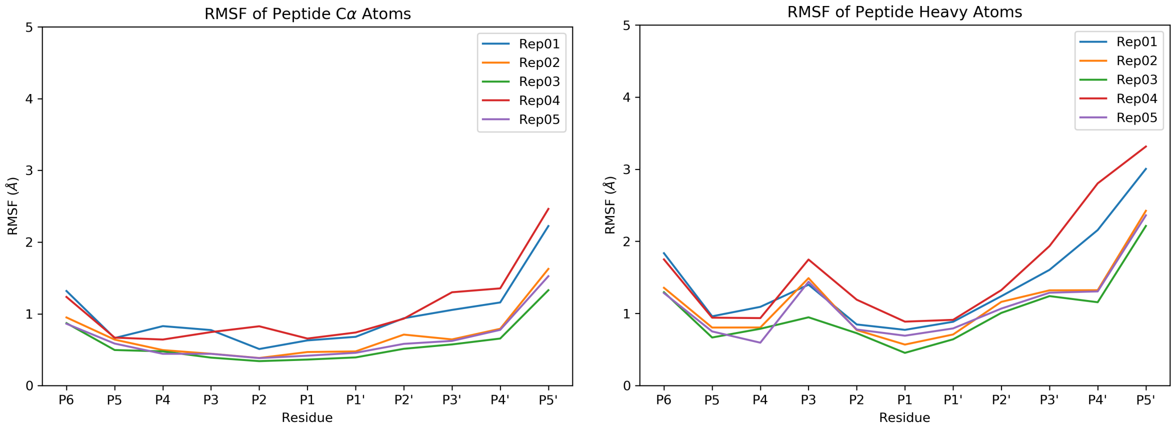


**Figure S1.7**: Per-residue RMSF of the s05 substrate peptide (left) Cα atoms and (right) heavy atoms in the 5 × 150 ns equilibrium MD simulations.

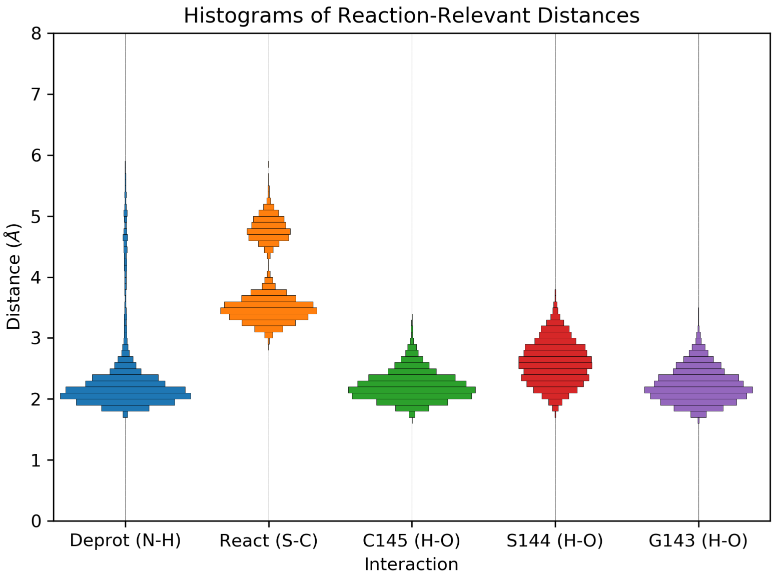


**Figure S1.8**: Distributions of selected distances relevant to the initiation of reaction between M$^{pro}$ Cys145 and the scissile substrate amide bond, observed over the combined 5 × 150 ns equilibrium MD simulations. These include (blue) the His41-NE2 – Cys145-HG distance; (orange) the Cys145-SG – substrate P1-C distance; and the distances from the substrate scissile amide carbonyl P1-O to the backbone NH atoms of (green) Cys145, (red) Ser144, and (purple) Gly143, which together form the oxyanion hole.

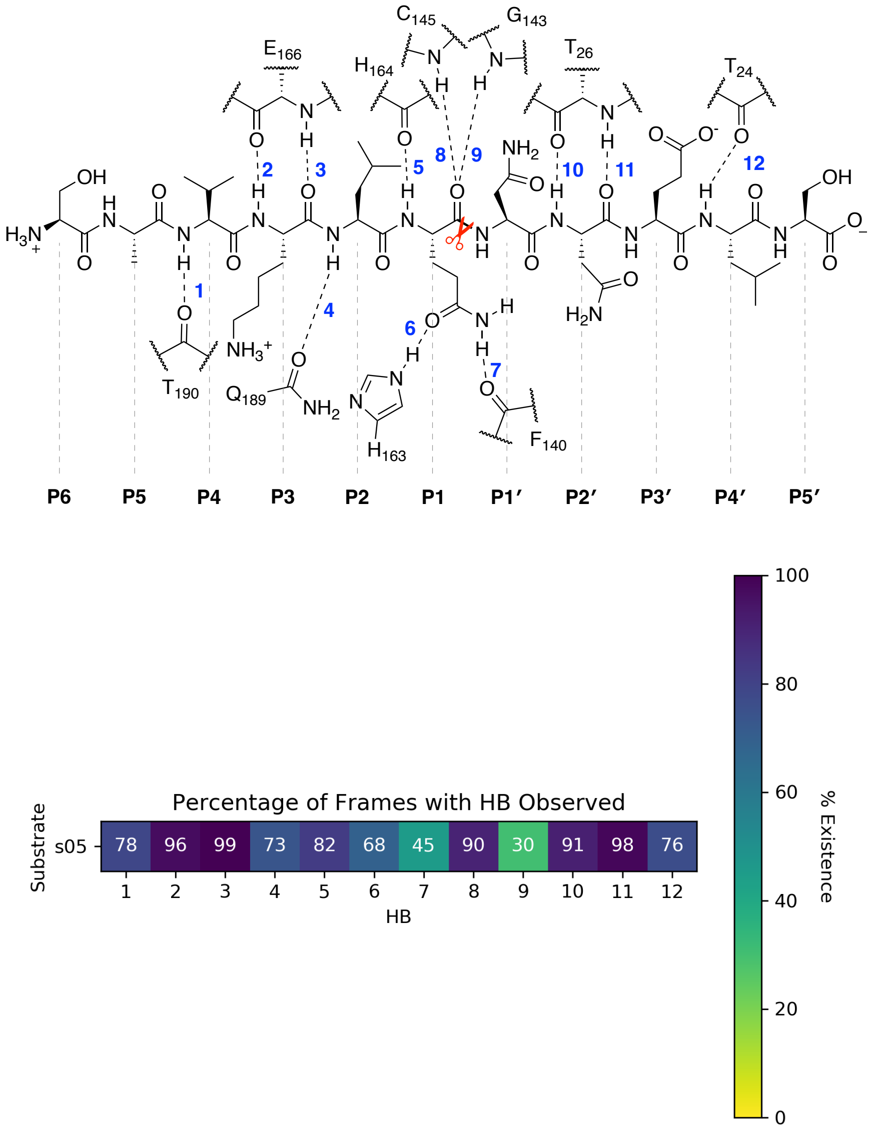


**Figure S1.9**: (top) Illustration of the 12 conserved M^pro^-substrate hydrogen bonds (HBs)[2] exemplified by s05; (bottom) a heatmap showing the percentage of frames (analysed every ns) where the HB is observed, over the combined 5 × 150 ns equilibrium MD simulations.

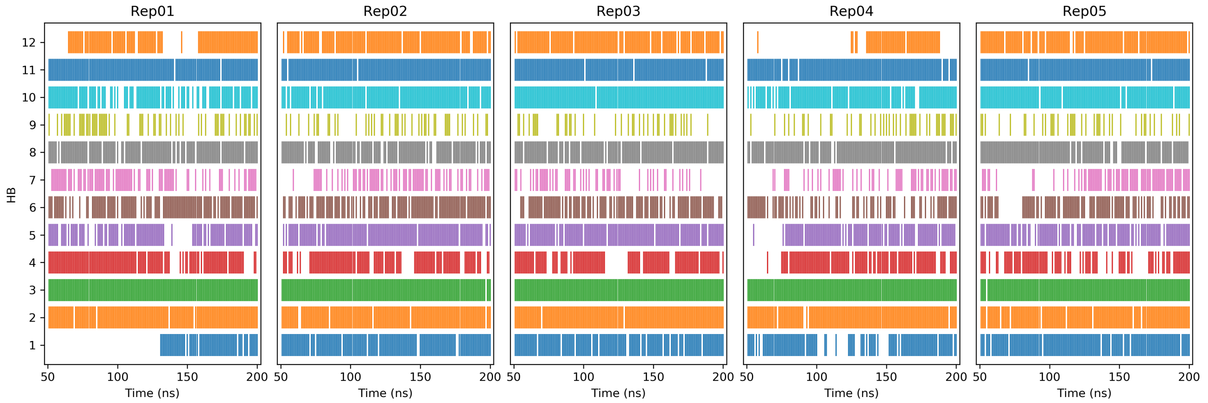


**Figure S1.10**: Event plot showing the existence of the 12 conserved HBs (analysed every ns) in each of the five 150 ns equilibrium MD simulations.

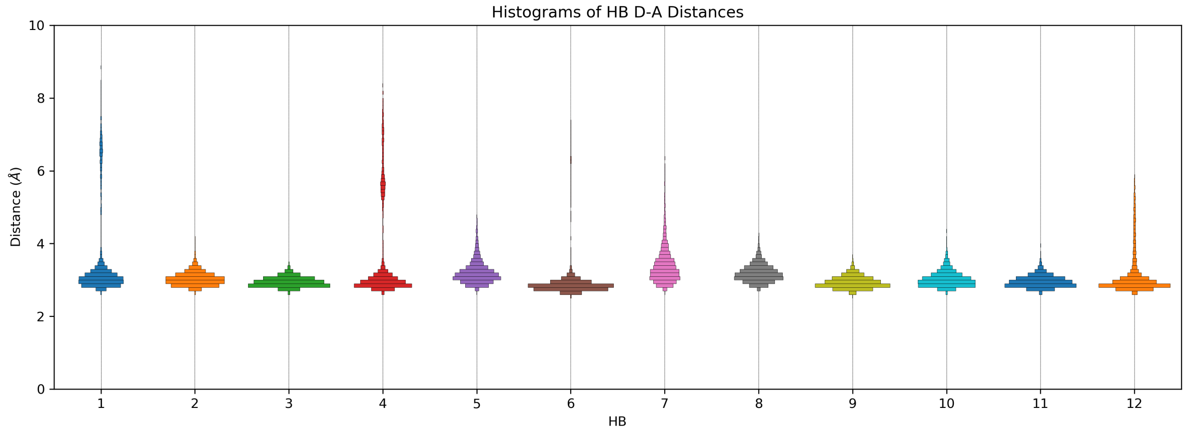


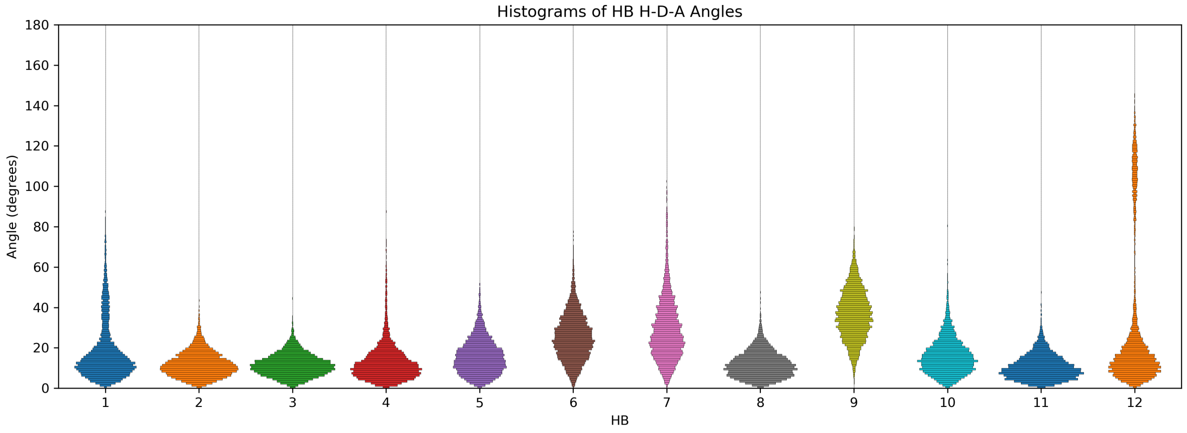


**Figure S1.11**: Distributions of (top) the donor-acceptor distances ($d_{D\text{-}A}$) and (bottom) the hydrogen-donor-acceptor angles (∠(H-D-A)) for each of the 12 conserved HBs, observed over the combined 5 × 150 ns equilibrium MD simulations. HBs are defined based on a combined distance ($d_{D\text{-}A} \leq 3.5$ Å) and angle (∠(H-D-A) ≤ 30°) criteria.

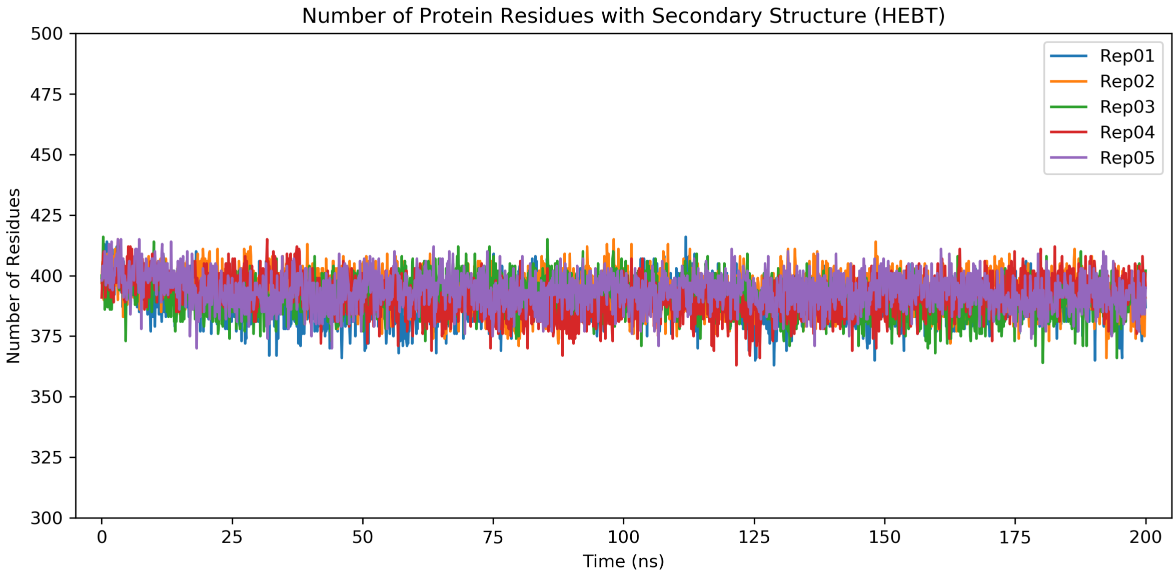


**Figure S1.12**: Time evolution of the number of M$^{pro}$ residues possessing secondary structure (H = α-helix; E = extended strand; B = β-bridge; T = hydrogen bonded turn) in each of the five 200 ns production MD simulations, as defined by DSSP (v 2.0.4).[3, 4]

### Section S2: Immediate effect of substrate removal

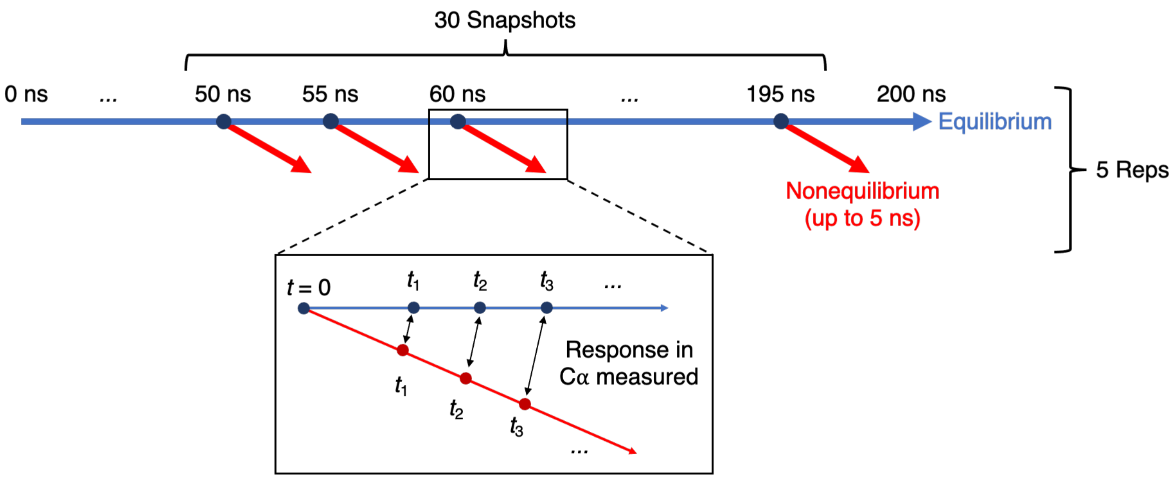


**Figure S2.1**: Scheme of the procedure used to set up and analyse the D-NEMD simulations. Five equilibrium MD simulations, each 200 ns, were performed for the M^pro-s05 substrate complex. These equilibrium simulations were used to generate starting structures for the nonequilibrium simulations. From the equilibrated part of each s05-bound simulation (from 50-200 ns), conformations were extracted every 5 ns, and the s05 peptide was removed from the system. The nonequilibrium simulations started from this ensemble of 150 conformations. The Kubo-Onsager approach was used to extract the response of M^pro to s05 peptide annihilation.[5, 6] For that, for each pair of equilibrium s05-bound and nonequilibrium apo simulations, the change in position for each Cα between the equilibrium and nonequilibrium trajectories at equivalent times was determined and averaged over all 150 simulations.

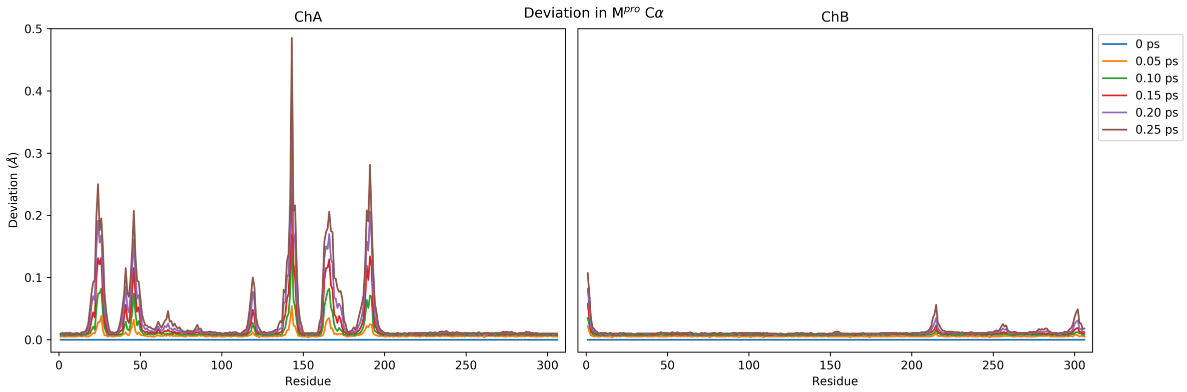


**Figure S2.2**: Cα deviations in (left) chain A and (right) chain B of the M^pro dimer as observed in the first six time points (saved in 0.05 ps intervals) up to 0.25 ps following the removal of s05.

**Table S2.1**: The five most perturbed M$^{pro}$ residues, in terms of Cα deviations, at selected time points within the 2 ps following the removal of s05.

| Rank | M$^{pro}$ residues with highest Cα deviations (Å) at each time point following perturbation | | | | | | | | | | | |
|---|---|---|---|---|---|---|---|---|---|---|---|---|
| | *t* = 0.10 ps | | *t* = 0.25 ps | | *t* = 0.50 ps | | *t* = 0.75 ps | | *t* = 1.00 ps | | *t* = 2.00 ps | |
| 1 | G143 | 0.168 | G143 | 0.485 | G143 | 0.623 | A191 | 0.706 | A191 | 0.813 | A191 | 1.132 |
| 2 | T26 | 0.082 | A191 | 0.281 | A191 | 0.547 | G143 | 0.693 | G143 | 0.781 | T190 | 0.905 |
| 3 | E166 | 0.082 | T24 | 0.250 | T24 | 0.448 | G23 | 0.603 | G23 | 0.684 | Q189 | 0.888 |
| 4 | M165 | 0.078 | N142 | 0.222 | N142 | 0.425 | T24 | 0.558 | T24 | 0.651 | T24 | 0.875 |
| 5 | T25 | 0.075 | C145 | 0.216 | G23 | 0.409 | N142 | 0.542 | S46 | 0.602 | S46 | 0.850 |

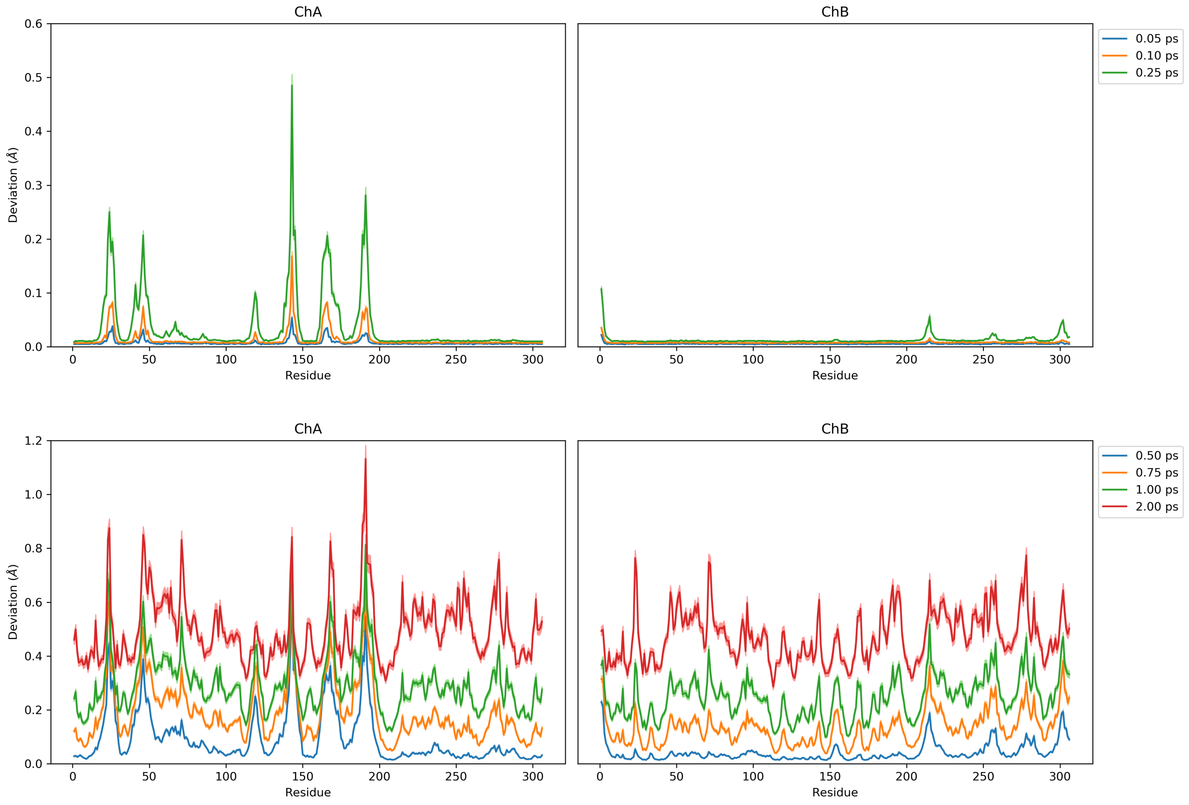


**Figure S2.3**: Cα deviations as observed up to 2 ps following perturbation (shown in two separate plots with different scales for clarity), with ± standard error of the mean (SEM) shown as shaded regions.

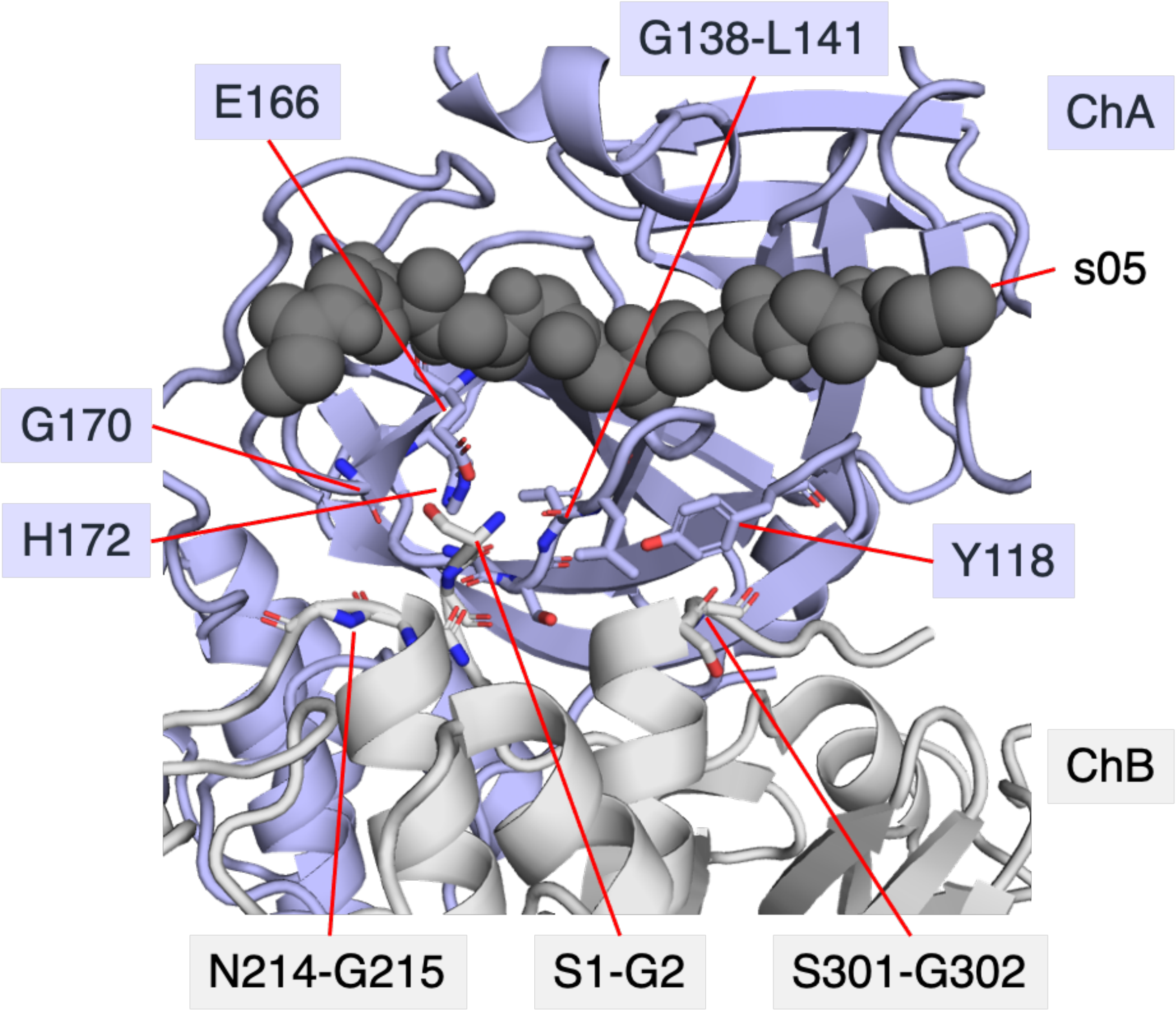


**Figure S2.4**: View of the dimer interface near the s05 binding site in the M$^{pro}$-s05 complex prior to MD simulations, with chains A and B shown in blue and grey, respectively, and the s05 peptide backbone shown as black spheres. Shown as sticks are chain B residues displaying substantial initial responses to s05 removal (deviation >15 pm at *t* = 0.50 ps), as well as their surrounding (within 3 Å) chain A residues, with labels in the corresponding chain colours.

### Section S3: Propagation of signal to allosteric sites

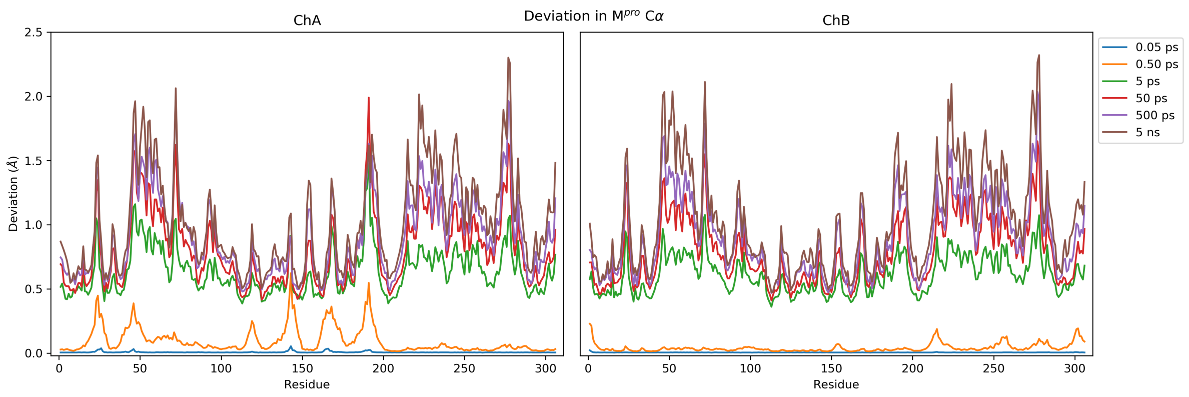


**Figure S3.1**: Cα deviations over the full range of the sampled timescales.

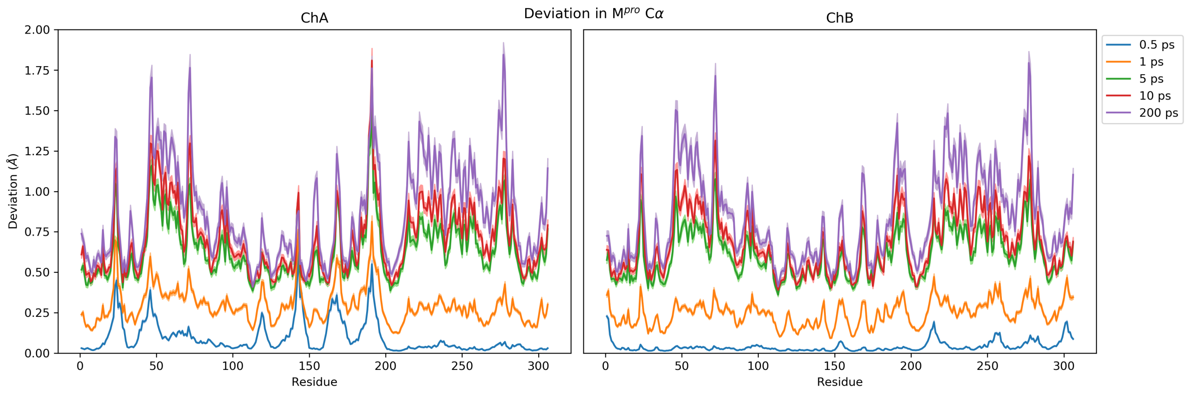


**Figure S3.2**: Cα deviations up to 200 ps, with ± SEM shown as shaded regions.

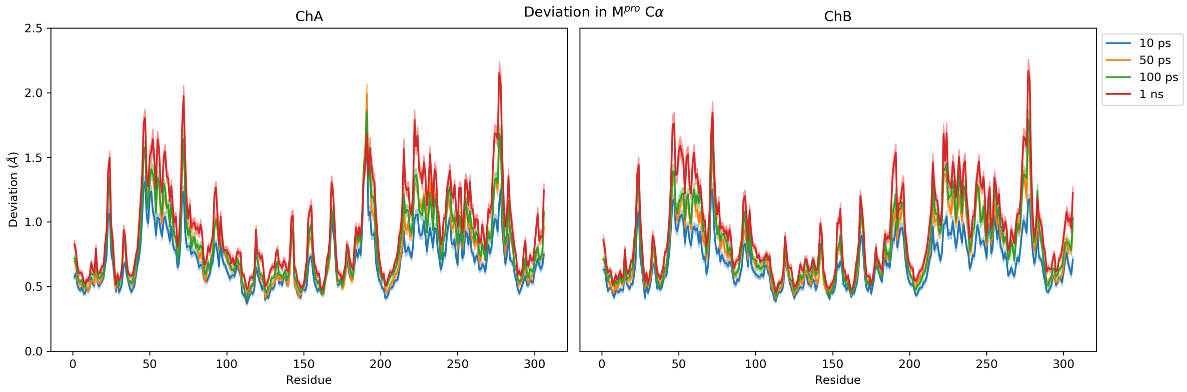


**Figure S3.3**: Cα deviations up to 1 ns, with ± SEM shown as shaded regions.

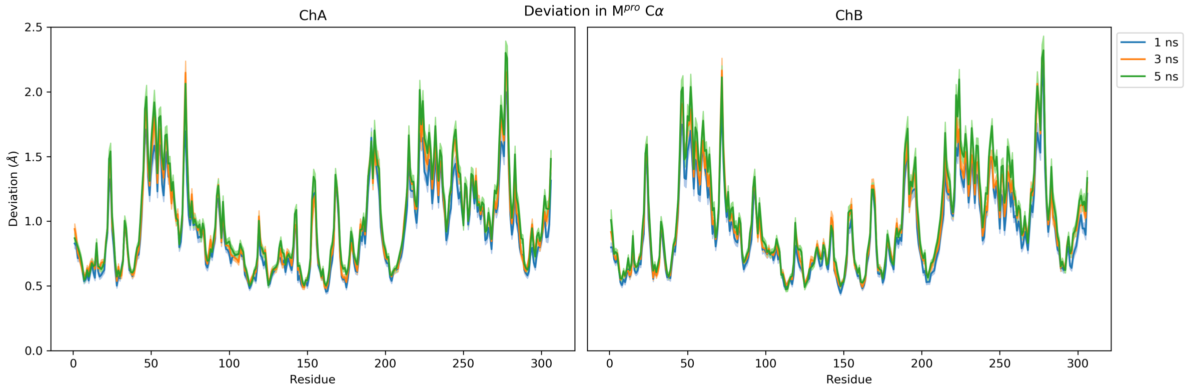


**Figure S3.4**: Cα deviations up to 5 ns, with ± SEM shown as shaded regions.

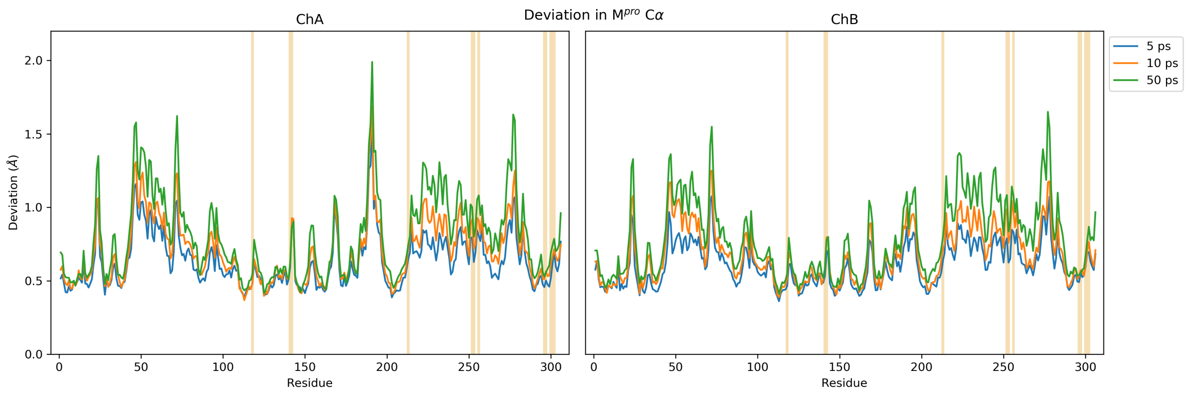


**Figure S3.5**: Cα deviations, with residues constituting the pelitinib binding allosteric site in both chains A and B (see **Figure 2** in main text) highlighted in light orange.

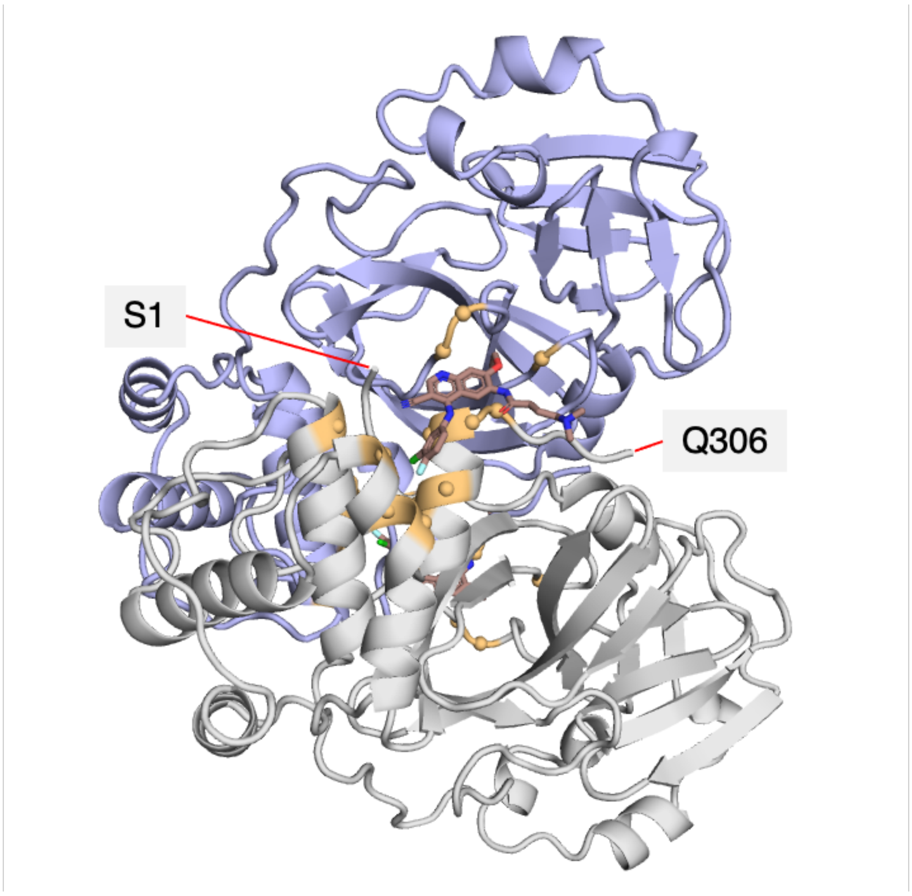


**Figure S3.6**: View of the $M^{pro}$ dimer (ChA and ChB shown in blue and grey respectively, with the N- and C-terminal residues of ChB labelled) complexed with two molecules of the allosterically binding inhibitor pelitinib (shown as brown sticks) (PDB 7AXM).[7] Residues constituting the allosteric sites (within 4 Å of pelitinib; see **Figure 2** in main text) are shown in orange.

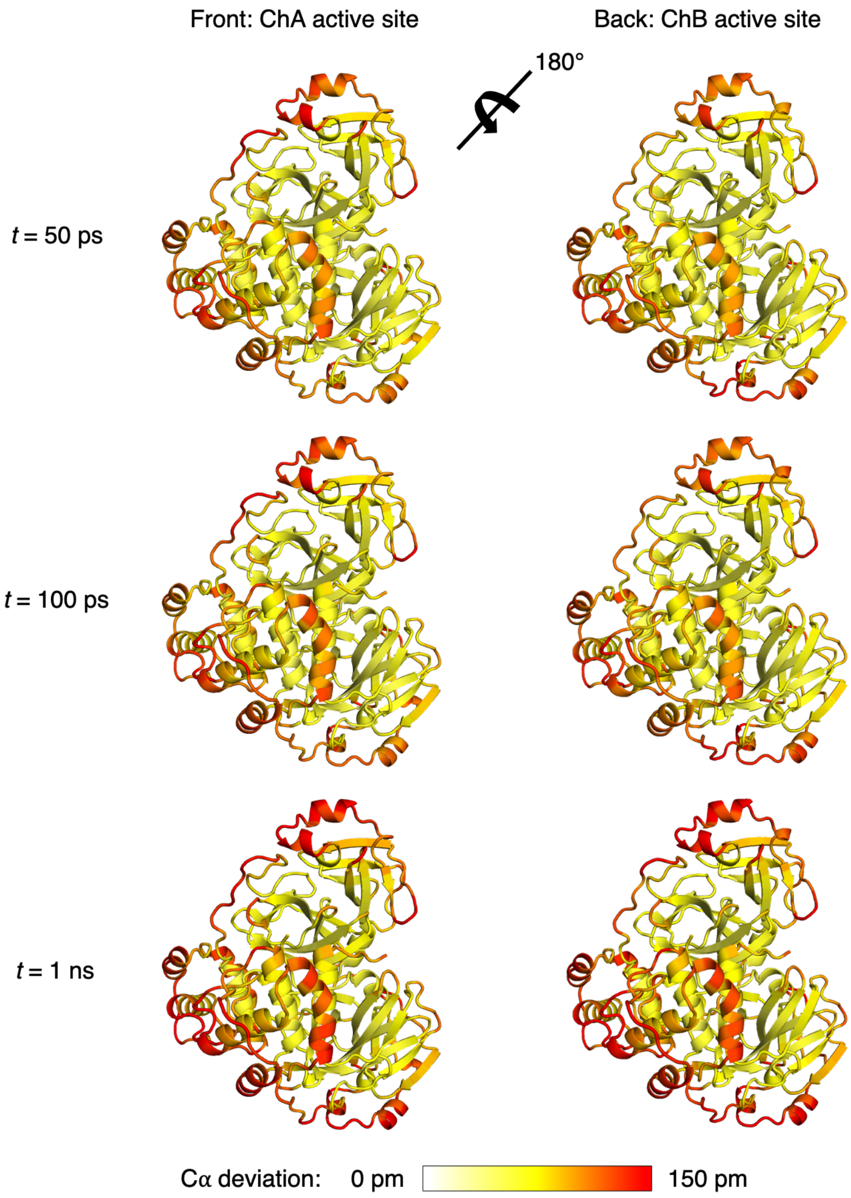


**Figure S3.7**: 180° rotated views of the M$^{pro}$-s05 complex prior to MD simulations, and the response to s05 removal in terms of average Cα deviations at *t* = 50 ps, 100 ps, and 1 ns, represented on a white-yellow-red scale. All figures were generated with PyMOL.[8]

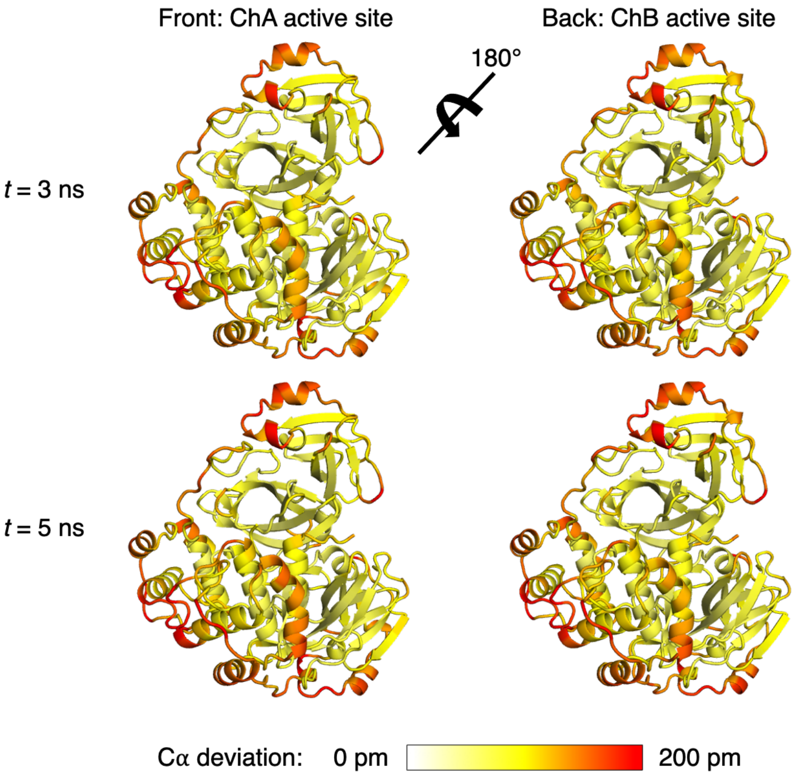


**Figure S3.8**: 180° rotated views of the M$^{pro}$-s05 complex prior to MD simulations, and the response to s05 removal in terms of average Cα deviations beyond 1 ns, represented on a white-yellow-red scale. Note the different scale used here compared to the previous figure to illustrate more clearly the intensified responses. All figures were generated with PyMOL.[8]

### Section S4: Coherent displacements in residues upon substrate removal

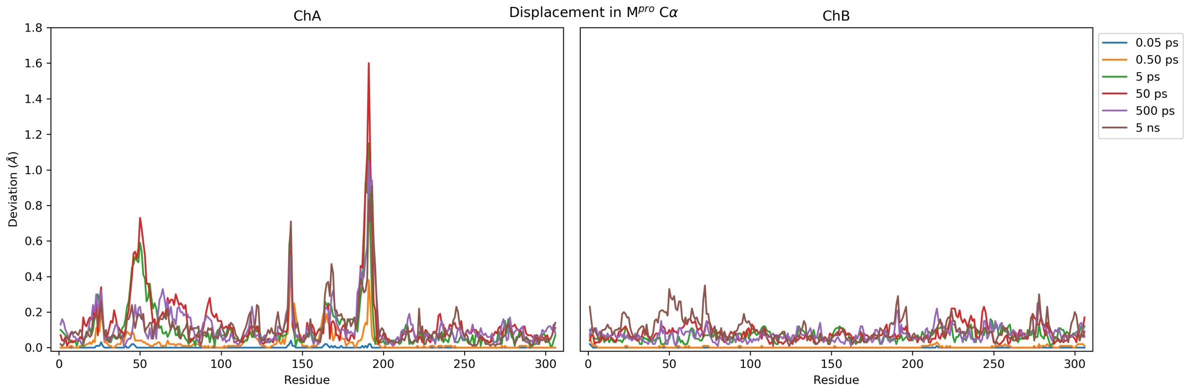


**Figure S4.1**: Magnitudes of average Cα displacement vectors over the full range of sampled timescales.

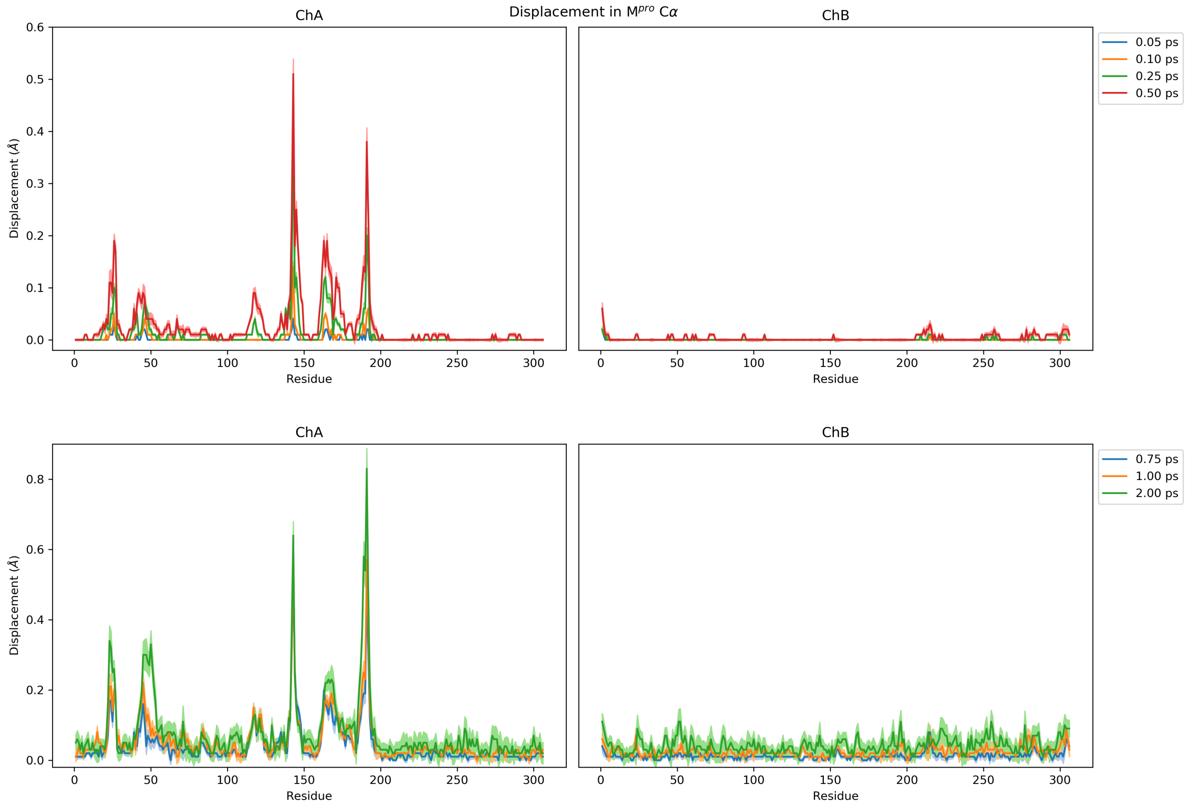


**Figure S4.2**: Magnitudes of average Cα displacement vectors (shown in two separate plots with different scales for clarity) up to 2 ps, with ± SEM shown as shaded regions.

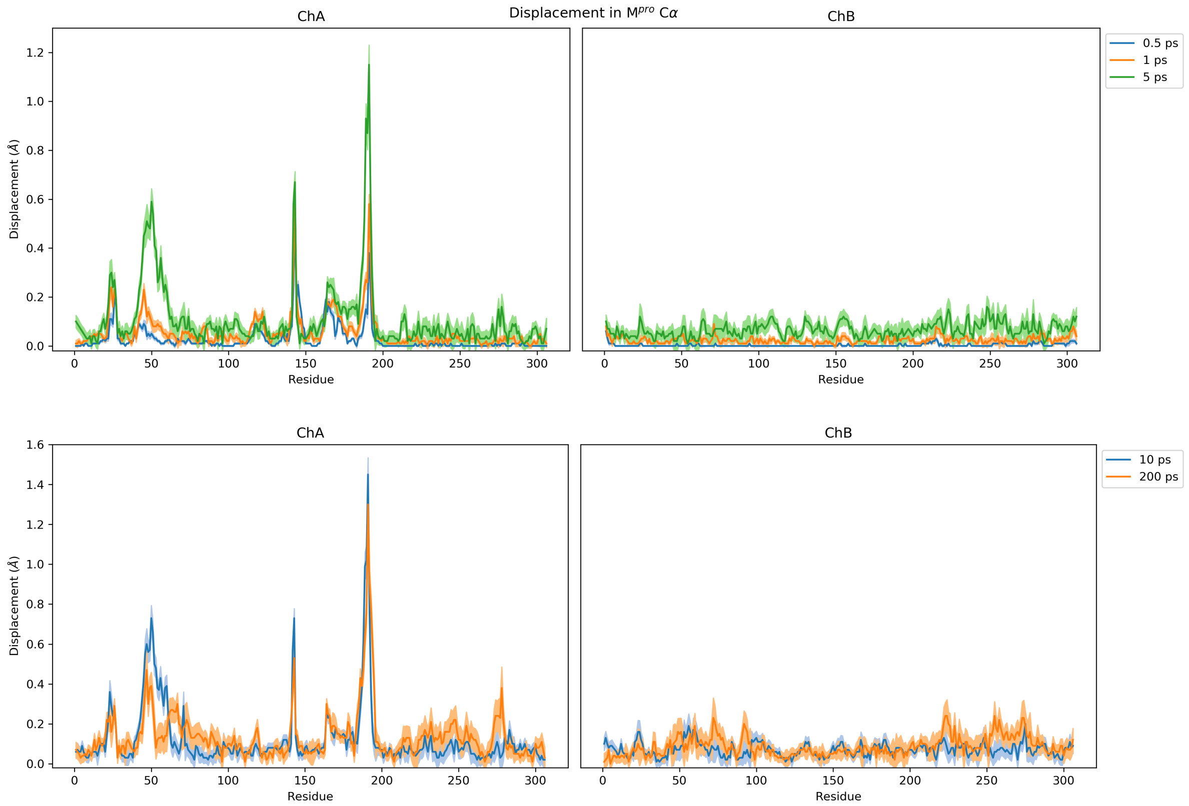


**Figure S4.3**: Magnitudes of average Cα displacement vectors (shown in two separate plots with different scales for clarity) up to 200 ps, with ± SEM shown as shaded regions.

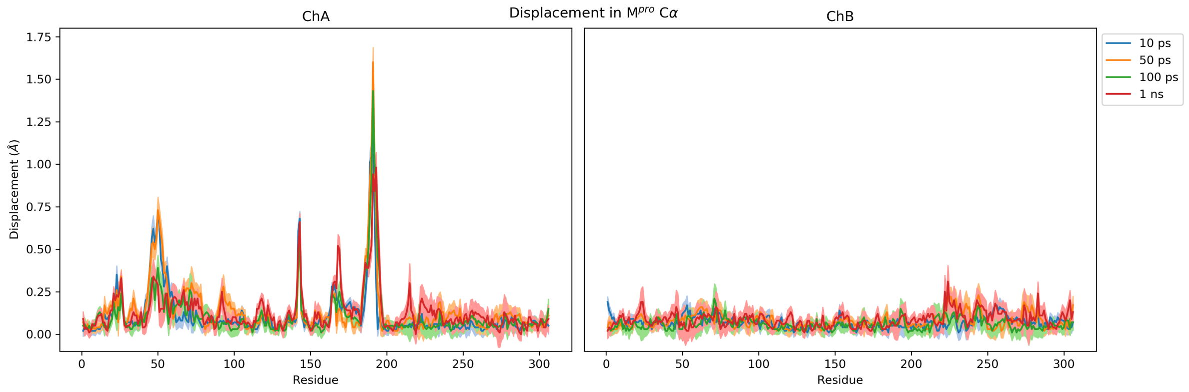


**Figure S4.4**: Magnitudes of average Cα displacement vectors up to 1 ns, with ± SEM shown as shaded regions.

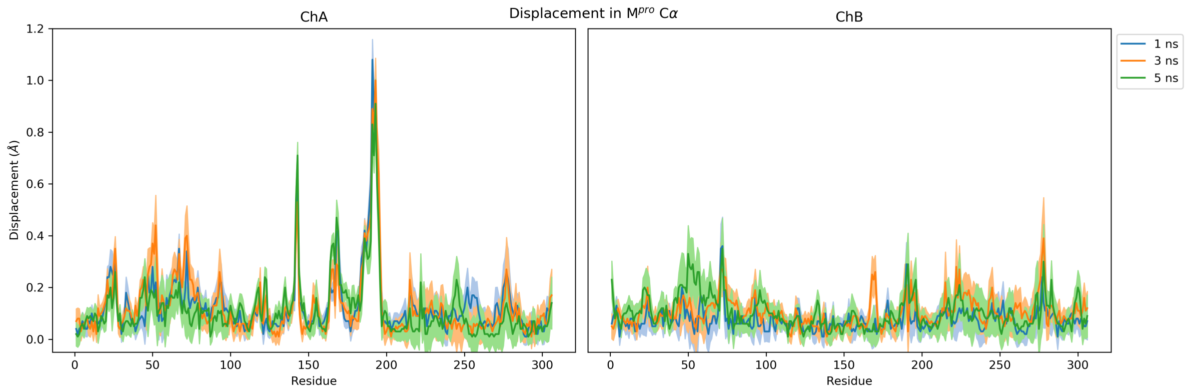


**Figure S4.5**: Magnitudes of average Cα displacement vectors up to 5 ns, with ± SEM shown as shaded regions.

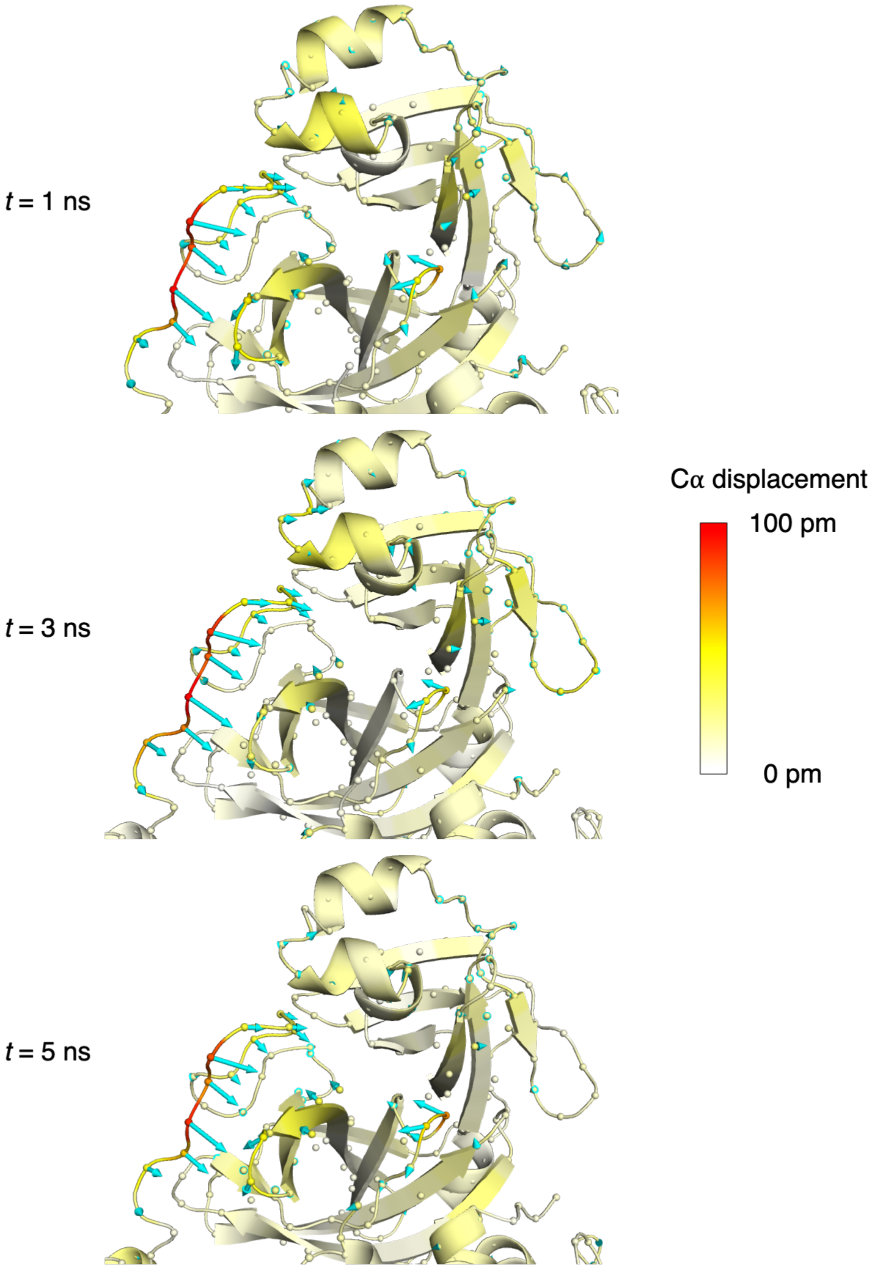


**Figure S4.6**: Views of the M$^{pro}$-s05 complex prior to MD simulations with a focus around the original substrate binding site, and the M$^{pro}$ response to s05 removal beyond 1 ns from averaging Cα displacement vectors. Displacement magnitudes are represented on a white-yellow-red scale.[8] Vectors with length ≥15 pm are displayed as cyan arrows with a scale-up factor of 5.[9]

**Table S5.1**: Protonation states of the histidine residues of SARS-CoV-2 $M^{pro}$ (PDB 6YB7)[1] in this study. The setup is identical to that reported in a previous study.[2] Following standard AMBER force field nomenclature,[10] neutral histidine residues that are N$\delta$-protonated and N$\epsilon$-protonated are referred to as "HID" and "HIE" respectively.

| His | State |
|---|---|
| 41 | HID |
| 64 | HIE |
| 80 | HID |
| 163 | HIE |
| 164 | HIE |
| 172 | HIE |
| 246 | HIE |

### References


1. Owen, C. D.; Lukacik, P.; Strain-Damerell, C. M.; Douangamath, A.; Powell, A. J.; Fearon, D.; Brandao-Neto, J.; Crawshaw, A. D.; Aragao, D.; Williams, M.; Flaig, R.; Hall, D.; McAauley, K.; Stuart, D. I.; von Delft, F.; Walsh, M. A. COVID-19 main protease with unliganded active site. *To be published* **2020,** *PDB 6YB7*, doi: 10.2210/pdb6yb7/pdb.
2. Chan, H. T. H.; Moesser, M. A.; Walters, R. K.; Malla, T. R.; Twidale, R. M.; John, T.; Deeks, H. M.; Johnston-Wood, T.; Mikhailov, V.; Sessions, R. B.; Dawson, W.; Salah, E.; Lukacik, P.; Strain-Damerell, C.; Owen, C. D.; Nakajima, T.; Świderek, K.; Lodola, A.; Moliner, V.; Glowacki, D. R.; Spencer, J.; Walsh, M. A.; Schofield, C. J.; Genovese, L.; Shoemark, D. K.; Mulholland, A. J.; Duarte, F.; Morris, G. M. Discovery of SARS-CoV-2 Mpro peptide inhibitors from modelling substrate and ligand binding. *Chem. Sci.* **2021,** *12* (41), 13686-13703.
3. Touw, W. G.; Baakman, C.; Black, J.; te Beek, T. A. H.; Krieger, E.; Joosten, R. P.; Vriend, G. A series of PDB-related databanks for everyday needs. *Nucleic Acids Res.* **2015,** *43* (Database issue), D364-D368.
4. Kabsch, W.; Sander, C. Dictionary of protein secondary structure: Pattern recognition of hydrogen-bonded and geometrical features. *Biopolymers* **1983,** *22* (12), 2577-2637.
5. Ciccotti, G.; Ferrario, M. Non-equilibrium by molecular dynamics: a dynamical approach. *Mol. Simul.* **2016,** *42* (16), 1385-1400.
6. Oliveira, A. S. F.; Ciccotti, G.; Haider, S.; Mulholland, A. J. Dynamical nonequilibrium molecular dynamics reveals the structural basis for allostery and signal propagation in biomolecular systems. *Eur. Phys. J. B* **2021,** *94* (7), 144.
7. Günther, S.; Reinke, P. Y. A.; Fernández-García, Y.; Lieske, J.; Lane, T. J.; Ginn, H. M.; Koua, F. H. M.; Ehrt, C.; Ewert, W.; Oberthuer, D.; Yefanov, O.; Meier, S.; Lorenzen, K.; Krichel, B.; Kopicki, J.-D.; Gelisio, L.; Brehm, W.; Dunkel, I.; Seychell, B.; Gieseler, H.; Norton-Baker, B.; Escudero-Pérez, B.; Domaracky, M.; Saouane, S.; Tolstikova, A.; White, T. A.; Hänle, A.; Groessler, M.; Fleckenstein, H.; Trost, F.; Galchenkova, M.; Gevorkov, Y.; Li, C.; Awel, S.; Peck, A.; Barthelmess, M.; Schlünzen, F.; Lourdu Xavier, P.; Werner, N.; Andaleeb, H.; Ullah, N.; Falke, S.; Srinivasan, V.; França, B. A.; Schwinzer, M.; Brognaro, H.; Rogers, C.; Melo, D.; Zaitseva-Doyle, J. J.; Knoska, J.; Peña-Murillo, G. E.; Mashhour, A. R.; Hennicke, V.; Fischer, P.; Hakanpää, J.; Meyer, J.; Gribbon, P.; Ellinger, B.; Kuzikov, M.; Wolf, M.; Beccari, A. R.; Bourenkov, G.; von Stetten, D.; Pompidor, G.; Bento, I.; Panneerselvam, S.; Karpics, I.; Schneider, T. R.; Garcia-Alai, M. M.; Niebling, S.; Günther, C.; Schmidt, C.; Schubert, R.; Han, H.; Boger, J.; Monteiro, D. C. F.; Zhang, L.; Sun, X.; Pletzer-Zelgert, J.; Wollenhaupt, J.; Feiler, C. G.; Weiss, M. S.; Schulz, E.-C.; Mehrabi, P.; Karničar, K.; Usenik, A.; Loboda, J.; Tidow, H.; Chari, A.; Hilgenfeld, R.; Uetrecht, C.; Cox, R.; Zaliani, A.; Beck, T.; Rarey, M.; Günther, S.; Turk, D.; Hinrichs, W.; Chapman, H. N.; Pearson, A. R.; Betzel, C.; Meents, A. X-ray screening identifies active site and allosteric inhibitors of SARS-CoV-2 main protease. *Science* **2021,** *372* (6542), 642-646.
8. Schrödinger LLC. *The PyMOL Molecular Graphics System, Version 2.3.0*.
9. Law, S. M. *modevectors.py*, https://pymolwiki.org/index.php/Modevectors, 2012 (accessed 2022-06-30).
10. Lindorff-Larsen, K.; Piana, S.; Palmo, K.; Maragakis, P.; Klepeis, J. L.; Dror, R. O.; Shaw, D. E. Improved side-chain torsion potentials for the Amber ff99SB protein force field. *Proteins* **2010,** *78* (8), 1950-1958.
